# Clock Work: Deconstructing the Epigenetic Clock Signals in Aging, Disease, and Reprogramming

**DOI:** 10.1101/2022.02.13.480245

**Authors:** Morgan E Levine, Albert Higgins-Chen, Kyra Thrush, Christopher Minteer, Peter Niimi

## Abstract

Epigenetic clocks have come to be regarded as powerful tools for estimating aging. However, a major drawback in their application is our lack of mechanistic understanding. We hypothesize that uncovering the underlying biology is difficult due to the fact that epigenetic clocks are multifactorial composites: They are comprised of disparate parts, each with their own causal mechanism and functional consequences. Thus, only by deconstructing epigenetic clock signals will it be possible to glean biological insight. Here we clustered 5,717 clock CpGs into twelve distinct modules using multi-tissue and in-vitro datasets. We show that epigenetic clocks are comprised of different proportions of modules, which may explain their discordance when simultaneously applied in a given study. We also observe that epigenetic reprogramming does not ‘reset’ the entire clock and instead the observed rejuvenation is driven by a subset of modules. Overall, two modules stand-out in terms of their unique features. The first is one of the most responsive to epigenetic reprogramming; is the strongest predictor of all-cause mortality; and shows increases with in vitro passaging up until senescence burden begins to emerge. The light-second module is moderately responsive to reprogramming; is very accelerated in tumor vs. normal tissues; and tracks with passaging in vitro even as population doublings decelerate. Overall, we show that clock deconstruction can identify unique DNAm alterations and facilitate our mechanistic understanding of epigenetic clocks.

## INTRODUCTION

Across every organ system—from cardiovascular to neurological—age stands out as the single biggest driver of disease^1^. Though the connection between risk and time may appear probabilistic on the surface, the emerging pathology is rooted in the molecular and cellular remodeling of the organismal system over its lifetime. Such changes likely result from accumulated damage, selection pressures at the level of cells, compensatory mechanisms, and/or the unintended consequences of a biological program. However, alterations to a complex system must abide by a hierarchical structure^2^, initiating at lower levels of biological organization (e.g. molecules) prior to manifesting at the higher levels in which they are typically observed (e.g. tissue and organ dysregulation and failure, and eventually death)^3^. Thus, to delay, prevent, or even reverse the maladies currently awaiting us in late life, we must discover how to decipher and remodel the molecular fingerprint of aging.

Alterations to the epigenome are one of the central molecular hallmarks of aging^4^, with potentially vast consequences for the physical and functional characteristics of cells. While a cell’s genetic code is essentially fixed, the epigenome is a dynamic master conductor, directing information encoded in DNA to generate the diversity of cells and tissues^5^. In many ways, it is akin to the ‘operating system of a cell’, controlling cell turnover rate, propagating cellular stress response, and supporting the maintenance and stability of cell populations in tissues. Unfortunately, the epigenetic program is also rewired over the lifespan^6^, leading some to hypothesize that epigenetic change may be the root source of aging-related phenotypes^7^.

One of the most extensively studied epigenetic aging phenomena is the alteration in the pattern of DNA methylation (DNAm). In mammals, DNAm refers to the covalent bond between a methyl group (CH3) and a CpG dinucleotide (5’—C—phosphate—G—3’)^8^. The presence of DNAm at CpG sites reduces regional accessibility due to folding and condensement of the DNA molecule around hydrophobic methyl groups^9^. Thus, gains in local DNAm are thought to be repressive and may explain preferential silencing of genes with age, while demethylation of CpGs may account for the age-associated loss of heterochromatin and unwanted activation transposable elements. Starting in 2011, DNAm patterns were found to be systematic to a degree that enable their use for developing ‘clocks’ aimed at estimating aging in cells and tissues^10^. This field has since boomed because of affordable array-based technologies and massive amounts of publicly available human methylation data. To date, there are more than a dozen such epigenetic clocks being applied to answer questions about aging, disease risk, and determinants of health^11^. Overall, epigenetic clocks have been shown to strongly track with age across a vast array of tissue and cell types^12^—even when trained using only data from blood^13^. More importantly, the discordance between predicted age and observed age appears to hold biological meaning. Increased epigenetic age relative to chronological age has been linked to disease phenotypes, mortality risk, and/or deleterious health behaviors^13–22^. Recently, much of the focus on epigenetic clocks has shifted towards examining them in the context of cellular reprogramming. Intriguingly, the conversion of somatic cells into induced pluripotent stem cells (iPSCs) via expression of Yamanaka factors can reverse the epigenetic aging signal—taking cells all the way back to a predicted age of around zero ^12,23–26^. However, it remains to be shown to what extent this truly represents an aging rejuvenation event. It is also unclear whether all DNAm age changes that accumulated within a cell are reversed, and if not, what the specific relevance is for those that are, versus are not, “reprogrammed”.

This lack of insight stems from an overall deficiency in mechanistic understanding of the changes captured by epigenetic clocks—what initiates these epigenetic changes and how or why are they implicated in disease etiology? Moreover, the debate over whether they are causal drivers versus casual passengers of aging has yet to be settled. The major obstacle we observe in uncovering mechanistic understanding relates to the way epigenetic clocks have been constructed. Epigenetic clocks are composite variables developed from a top-down perspective that combines input from typically hundreds to thousands of CpGs that appear to change with aging, without regard to the underlying biology^27^. As such, they likely are comprised of many different subtypes of methylation patterns—each with its own causal explanations and functional consequences. For instance, gains in DNAm at CpG islands located in promoter regions of genes may differ in their mechanistic underpinnings as compared to CpGs representing loss of DNAm at intergenic enhancers. For this reason, insights into the causes and consequences of DNAm age changes will require a reductionist approach.

In this paper we combined computational and experimental approaches to deconstruct epigenetic clocks and group CpGs into smaller functionally related modules, from which epigenetic aging mechanisms can be more easily discovered. We demonstrate that not all signals captured in the clocks are equal when it comes to morbidity/mortality risk. We also show that reprogramming is concentrated on a few specific modules, yet the discrepancy in response across CpGs is not decipherable at the level of the whole clock.

## RESULTS

### Epigenetic clock CpGs can be grouped into characteristically distinct modules

A schematic of the methods used to deconstruct epigenetic clocks is presented in Figure 1. We started by compiling a list of multiple Illumina DNAm array datasets from human tissues and *in vitro* human cell lines (Table S1). We then restricted these datasets to 5,717 CpGs that are present in fifteen of the most well-known epigenetic clocks (Table S2), as well as five new tissue-specific clocks (blood, brain, liver, skin, and adipose) that we developed using only highly conserved (mouse-human) CpGs present on the human 450k array, EPIC array, and the new Mammalian Methylation Array. Overall, this resulted in seven data matrices each with 5,717 parameters (CpG beta-values) and between 13 and 2,478 observations. Next, we utilized a novel consensus spectral clustering method to group each of the 5,717 CpGs into a module. Details can be found in methods, but briefly, affinity matrices were constructed for each dataset using both Euclidean and biweight midcorrelation distances. These were then combined to form a single Meta-Affinity matrix by taking the sample quantile at probability p=0.25. This method identifies clusters of CpGs that both co-vary (e.g. change in a concerted manner with age) and share similar ranges of methylation values. Once the single affinity matrix was estimated, Spectral clustering was carried out using traditional methodology.

**Figure 1:**
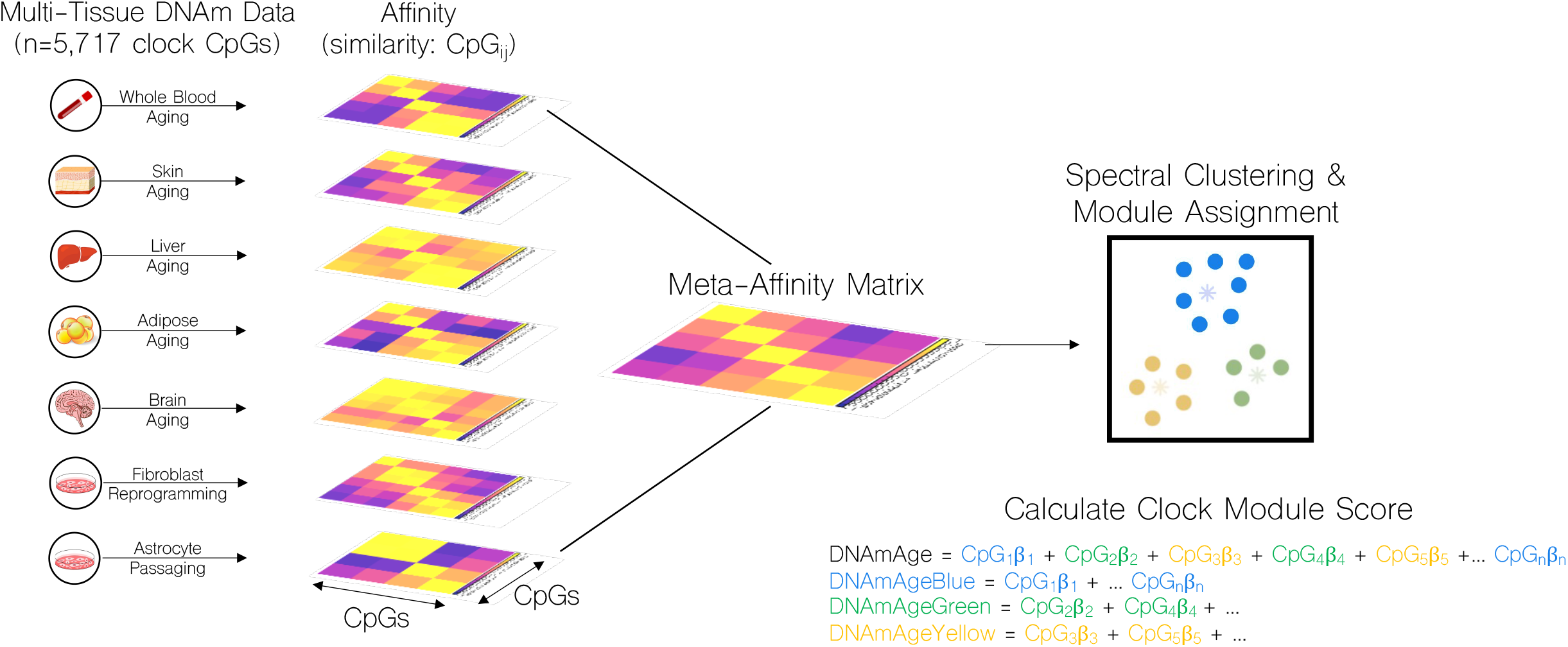
Schematic of epigenetic clock deconstruction. We compiled seven human DNAm datasets and restricted them to CpGs found in any one of fifteen commonly used epigenetic clocks (n=5,717 CpGs). For each dataset, we estimated relationships between CpG pairs using affinity matrices based on biweight midcorrelations. For two datasets (whole blood and fibroblast reprogramming), we also generated Euclidean based affinity matrices. These were then combined into a single meta-affinity matrix on which we performed Spectral Clustering. CpGs were assigned to a given module—denoted by color— based on clustering. Clusters were filtered to remove assignments for those with <100 members. Using the original epigenetic clock equations, we next calculated module specific clock scores by setting CpGs not belonging to a given module to zero. This produced a total of 156 module clock combinations (e.g. ‘HorvathPanTissue Blue’) that were examined in subsequent analyses. Not all modules were found in all clocks.

This method produced 12 modules, each denoted using a specific color to distinguish them. Modules ranged in size from n=242 CpGs (green module) to n=705 CpGs (red module). Approximately 16% of clock CpGs (n=938) were ‘unassigned’. As shown in Figure 2, the various modules are comprised of CpGs that exhibit different behaviors regarding their dynamic ranges during aging and reprogramming (GSE54848). Six modules stand out—green, green-yellow, lightblue, cyan, red, and yellow. For instance, both the green and the green-yellow modules lose DNAm with age in blood (Figure 2A), and gain DNAm upon reprogramming (Figure 2B). The distinction is that this seems to happen to a greater degree in the green-yellow module. In iPSCs, it is more hypermethylated than the green module (p<2e-16), but in adult blood, the two modules show more similar levels of DNAm. This may suggest that CpGs in the green-yellow are more likely to be methylated in pluripotent cells, but during the differentiation process, the two modules converge. We also observe differences in the locations of CpGs in these two modules. The green module is highly enriched for CpGs in Shores (regions with moderate CpG density located in proximally to high density regions), while the green-yellow module is enriched for CpGs in Open Seas (low density CpG regions).

**Figure 2:**
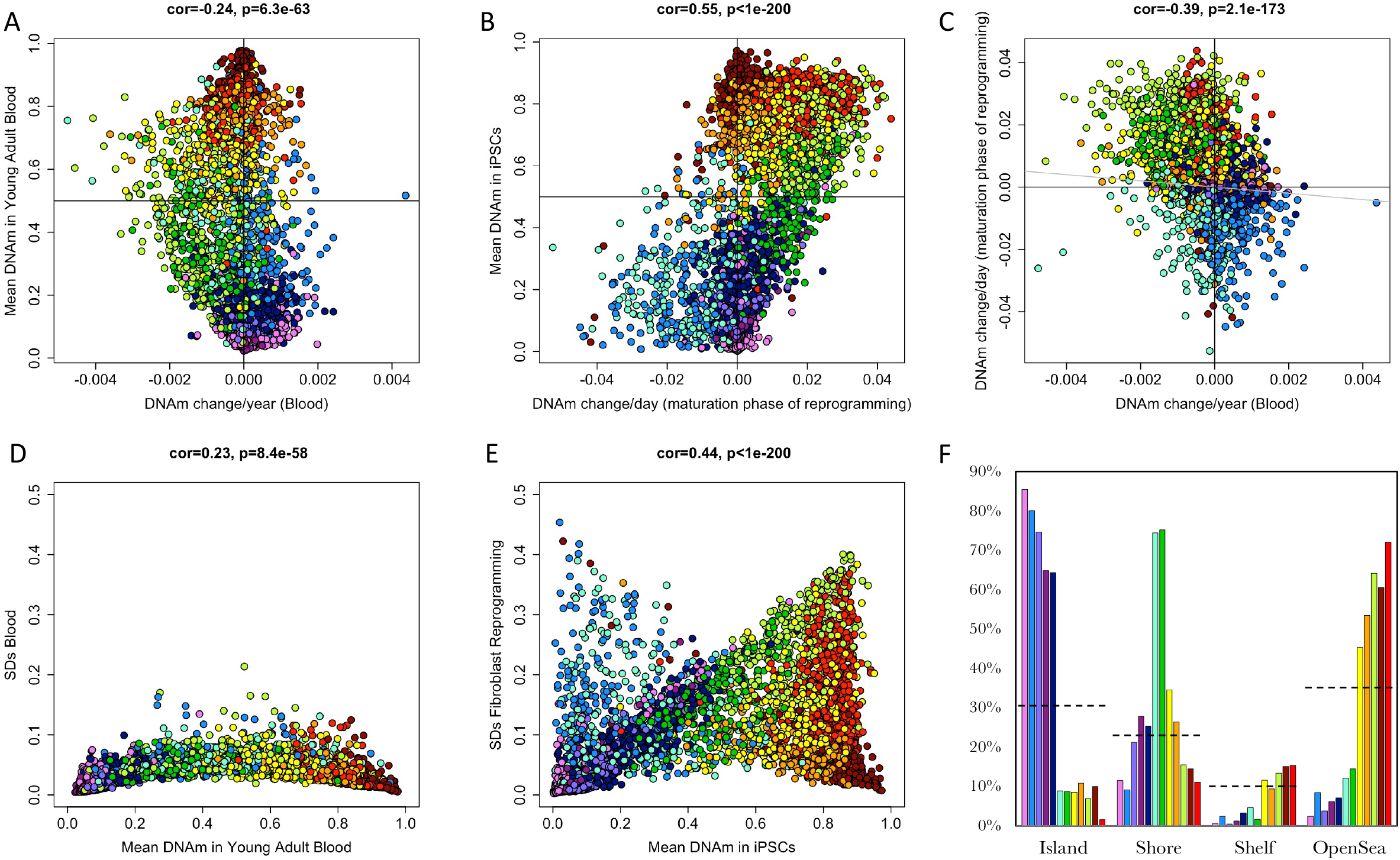
Descriptive characteristics of module CpGs. For each CpG, we estimated the degree of DNAm change per year in adults by fitting a linear regression model in the Framingham Heart Study (FHS). (A) Change per year in DNAm was plotted against the mean DNAm level for each CpG estimated in adults less than 35 years of age from FHS. Point colors denote module assignment, while the vertical line separates CpGs that increase versus decrease with age and the horizontal line distinguishes hypermethylation (DNAm>0.5) from hypomethylation (DNA<0.5). We observe a moderate to weak correlation, suggesting that CpGs with hypermethylation tend to lose DNAm with age, while those with hypomethylation gain DNAm with age. We also observe distinct module patterns, such that the green-yellow module and the light blue module have similar levels of DNAm in young adults, but they change in opposite directions. (B) We also plotted changes per day in DNAm during maturation phase (days 15-28) of epigenetic reprogramming via Yamanaka expression in dermal fibroblasts against mean DNAm in fully reprogrammed iPSCs. Again, point colors denote module assignment, while the vertical line separates CpGs that increase versus decrease with reprogramming and the horizontal line distinguishes hypermethylation (DNAm>0.5) from hypomethylation (DNA<0.5) in iPSCs. Here we observe a moderate positive correlation such that CpGs that are hypermethylated in iPSCs were the ones that gained the most DNAm during reprogramming and those that are hypomethylated in iPSCs were the ones that lost the most DNAm during reprogramming. We also observe modules with similar levels of DNAm in iPSCs, but with different degrees of change during reprogramming (e.g. orange and greenyellow). (C)We plotted change in DNAm with age against change in DNAm with reprogramming and observe a moderate negative correlation, suggesting that there is an association between the level of DNAm gain with age and the level of DNAm loss with reprogramming. (D) for each CpG, we plotted mean DNAm in young blood against variance (as measured by standard deviation) of DNAm in blood. (E) For each CpG, we plotted mean DNAm in iPSCs against variance (as measured by standard deviation) of DNAm during reprogramming. (F) Using annotation of CpGs, we find that modules vary in their relation to high CpG dense regions—CpG islands. The y-axis denotes the percent CpGs in a given module within each type of region, with different colored bars representing the 12 modules. Dashed horizontal lines indicate percent CpGs in a given region across all modules (n=5,717 CpGs).

The light-blue and cyan modules contain CpGs illustrating the other side of the spectrum. Both modules have CpGs that lose DNAm during reprogramming (Figure 2B). However, during aging, the light-blue module gains DNAm, while the cyan module loses DNAm (Figure 2A & 2C). This suggests a counterintuitive pattern for the cyan module such that aging and reprogramming (considered anti-aging) show similar direction of change in DNAm. Like the green module, we find that the cyan module is enriched for CpGs in Shores. The light-blue module, on the other hand, is enriched for CpGs in Islands (Figure 2F).

The red and yellow modules are similar to the green and green-yellow in that the CpGs assigned to them lose DNAm with age and regain DNAm with reprogramming. The difference is that this happens to a lesser extent, especially for age change in the red module (Figure 2A and Figure S1) and reprogramming changes in the yellow module (Figure 2B and 2C). The red module also tends to be more hypermethylated in both blood and iPSCs compared the green or greenyellow modules. When considering enrichment, the red module has the highest proportion of CpGs in Open Seas compared to any module (Figure 2C).

Finally, CpGs towards the two extremes (0 or 1), were assigned to either the dark-red (hypermethylation) or dark-magenta (hypomethylation) modules (Figure 2A and 2B). They tend to have very limited dynamic ranges and when it comes to a percent DNAm change, CpGs in these two modules are not altered much by either reprogramming or age (Fig 2D and 2E). When considering standardized changes (Figure S1), they do appear to be somewhat altered in aging and reprogramming, but not in a particular direction; that is, they do not systematically gain or lose DNAm as a function of age or reprogramming. One explanation is that we and others have reported that CpGs with mean levels near 0 or 1 tend to be plagued by technical noise to a much greater degree than CpGs that are not on the boundary (between 0.2 and 0.8). This is more likely than explanations relying upon randomization with aging: entropic contributions would bias the change as regression towards the mean (DNAm moving closer to 0.5). We also find that the dark-magenta module is enriched for CpGs in islands, while the dark-red module is enriched for CpGs in Open Seas. This is consistent with prior literature suggesting the CpG islands tend to be hypomethylated while Open Seas tend to be hypermethylated.

### Epigenetic clocks are comprised of differing proportions of CpGs from each module

Although epigenetic clocks were all developed to proxy the same biological phenomenon of aging, recent papers have shown that they often diverge when it comes to their associations with a variety of outcomes. Some clocks are better able to predict mortality from blood^19^; others can differentiate tumor versus normal tissue^28^; and still others can track perturbations induced in vitro^28,29^. While one explanation may be that this is due to the degree of error in each clock, we hypothesized instead that—given that epigenetic clocks are multifactorial composites—each clock may contain a different proportion of CpGs from a given module. Or moreover, different epigenetic mechanisms may be contributing more-or-less to the overall signals in various epigenetic clocks. To test this, we recalculated clocks scores, but on a module-by-module basis. For instance, most clocks are calculated as a weighted sum of CpGs (Figure 1). Thus, if we only sum the weighted CpGs from a given module, we get the portion of the clock signal that each modules contributing to. If you sum across all modules in a clock (including unassigned CpGs), you once again get the entire clock score.

We used data from the whole blood and reprogramming experiment used for Figure 2 to compare module proportions across clocks. These two datasets were compared because CpGs have different ranges of DNAm across tissue/cell types and thus different modules may be amplified or diminished in certain contexts. However, we find that the proportion of the overall clock signal that each module contributes to is mostly similar when comparing blood and reprogramming, with a few exceptions (Figure 3). The yellow module contributes to more of the signal in reprogramming versus blood for six clocks (Pan-Tissue^12^, PhenoAge^13^, SkinBlood^29^, DNAmTL^30^, Mammalian Blood and Mammalian Brain). Similarly, the green-yellow module contributes to more of the signal in the Pan-Tissue, PhenoAge, SkinBlood, Hannum^31^, and Mammalian Blood clocks. These gains in signal for the yellow and green-yellow modules tend to be counterbalanced by reductions in the proportion of signal coming from the light-blue and/or navy module in the context of fibroblast reprogramming versus blood aging.

**Figure 3:**
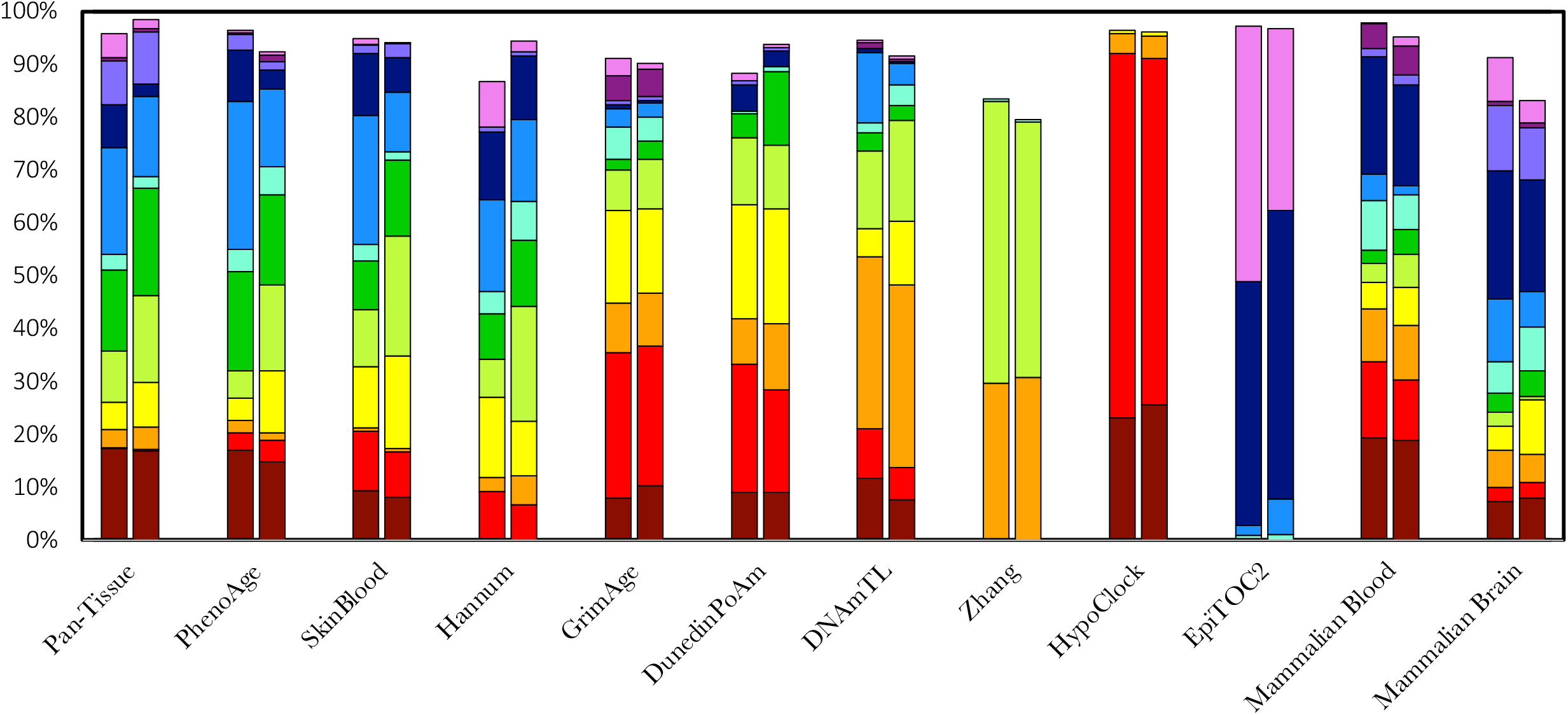
Module proportions across clocks. Module specific clocks were calculated for all clock-module combinations. Here we show the proportions of overall signal for twelve epigenetic clocks. The y-axis denotes the percent of the total sum of absolute values for each module. For each clock, the first stacked bar is the percentages estimated from whole blood and the second stacked bar is the percentages estimated in data from reprogramming of dermal fibroblasts. We observe differences between clocks when it comes to the degree of signal from each module. For instance, Pan-Tissue, PhenoAge, and SkinBlood clocks share similar module proportions, while GrimAge, DunedinPoAm, and DNAmTL share similar proportions. Three clocks—Zhang, HypoClock, and EpiTOC2—are comprised of predominantly two modules. Finally, all modules are observed in in mouse/human conserved mammalian clocks.

Perhaps more interesting and dramatic than the differences between blood and reprogramming within each clock, are the differences observed between the various clocks. For instance, the Pan-Tissue, PhenoAge, SkinBlood, and to some degree, the Hannum clock have similar makeups in terms of the proportions of their overall signal owed to each module. Conversely, GrimAge^19^ and DunedinPoAm^22^ share a very similar composition. DNAmTL is also similar to GrimAge and DunedinPoAm, with the exception of having an enriched signal from the orange module. Three clocks in particular are made of signals owing almost entirely to just two of the twelve modules. Zhang is mostly comprised of the orange and green-yellow module; HypoClock^32^ is mostly comprised of the dark-red and red modules; and EpiTOC2^33^ is mostly comprised of the navy and pink modules. Lastly, the two mammalian clocks do contain signals from all modules, suggesting that, at least to some degree, all modules are present in regions conserved between human and mouse. As the described composition similarities and contrasts of clocks reflect the already presumed similarities based on their trained uses, this may reflect some characterization of the underlying biological organization their training indirectly encoded.

### Differential age correlations between epigenetic clock modules

By calculating a module-specific score for each clock, we can test whether associations observed for the whole clocks can be attributed to specific modules. We started with the most robust associations attributed to the clocks—the correlation with chronological age. Specifically, the Horvath Pan-Tissue clock has been renowned for being able to predict chronological age across a wide array of tissues with very small mean squared error rate^12^. Therefore, we used data from whole blood, liver, brain, skin, and adipose to test if this tissue consistency remains the case across each of the modules that make up the Pan-Tissue clock. Additionally, the Pan-Tissue clock and the SkinBlood clock also include a logarithmic transformation for samples with ages less than 20 years^12,29^. This is used to account for the non-linear exponential increases in epigenetic age often observed during development. However, we excluded this step to test whether the non-linearity is in fact module specific.

As shown in Figure 4, modules vary in their multi-tissue age correlations, ranging from r=0.016 (pink) to r=0.76 (green-yellow). For the most part, the loss of robust age correlations can be attributed to one of two things—tissue differences, or non-linearity. For example, the dark-red, red, yellow, lilac, and purple modules show very striking differences in age prediction across tissues. However, they also differ in terms of which tissue they predict as ‘older’ vs. ‘younger’.

**Figure 4:**
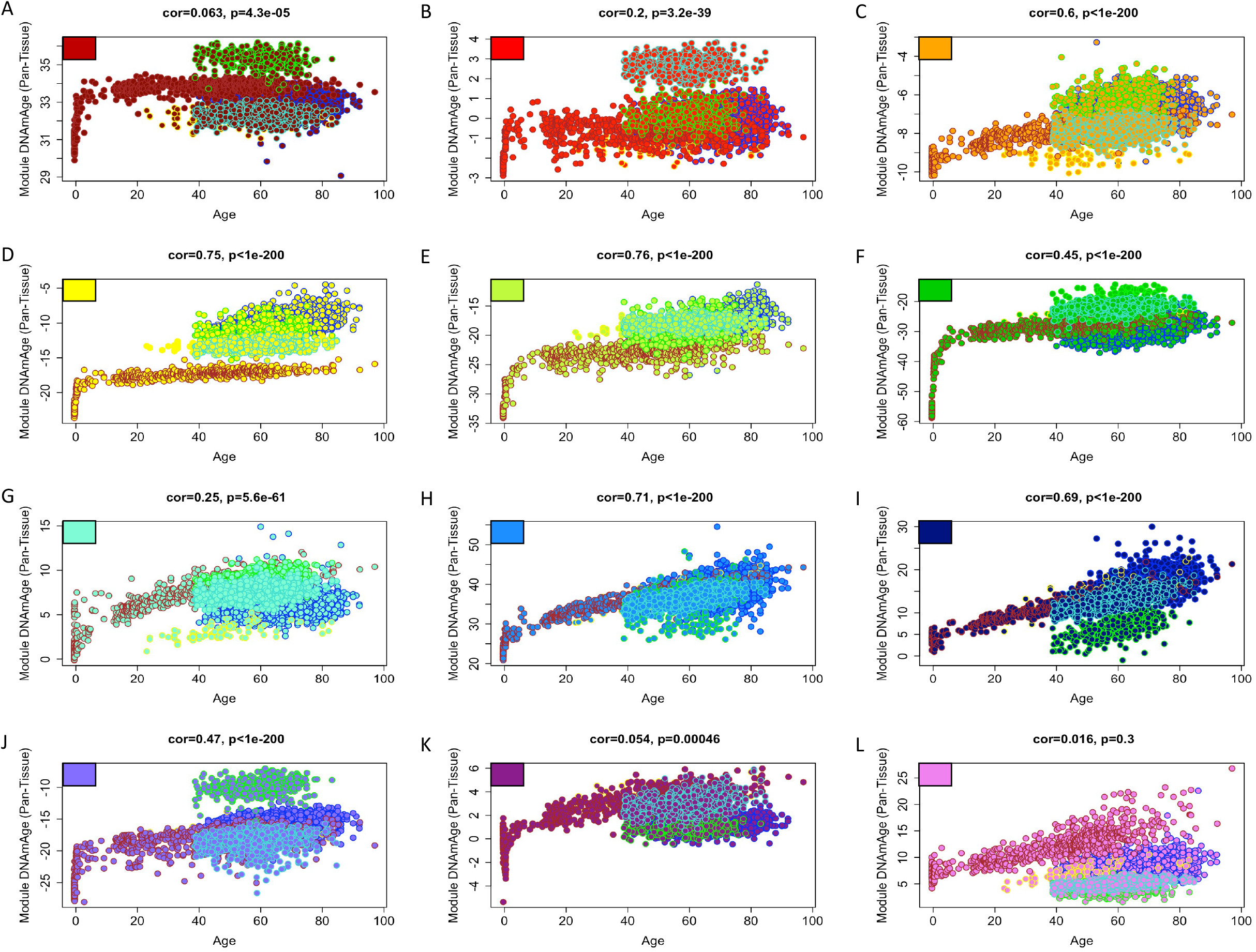
Multi-tissue age correlations of Pan-Tissue clock modules. Using the equation for the Horvath Pan-Tissue clock, we estimated module specific DNAmAges and plotted them (y-axis) against chronological age (x-axis) for blood, liver, skin brain, and adipose. Colors in the upper left and point colors represent the modules. Point outlines represent tissues—brain (brown), blood (blue), skin (green), adipose (turquoise), and liver (yellow). Despite the fact that the Pan-Tissue clock shows minimal tissue differences in age prediction, we observe tissue differences when examining at the module levels. We also observe differences between modules in the rate of change in DNAmAge during the developmental phase (only available for brain).

This likely averages out for the full clock score, producing consistent age predictions across tissues. We also notice that certain modules appear to increase exponentially during development, while others do not. The non-linear change appears most striking for the green and green-yellow modules, while the navy, orange, and pink modules are observed to be nearly linear.

We also considered tissue differences using modules for EpiTOC2 (Figure S4), given that this clock was designed to track cell divisions^33,34^ and thus more highly proliferative tissues should be found to be “older”. However, this was not necessarily the case across the modules. In the pink module brain was found to have higher values than blood, liver, skin, and adipose. For the navy module, the age-adjusted order was liver (highest), blood, brain, adipose, and skin; while for the light-blue module, liver was again the highest, followed by blood, skin, adipose, and finally brain. We also did not observe dramatic exponential increases in the module scores during growth and developmental periods, and instead, the EpiTOC2 modules appeared to increase somewhat linearly with age throughout the range.

HypoClock is another measure initially derived to track mitotic rates across cells and tissues^32^. Again, we did not observe any extreme acceleration or exponential increase during development for any of the modules (Figure S4). The HypoClock modules (the dark-red, red, and orange) instead showed only minimally accelerated changes during growth and development. When comparing tissues, the dark-red module estimated liver as oldest, then blood, brain, adipose, and skin. The red modules, ranked them: liver, blood, adipose, brain, and skin, which was almost the same for the orange module, except that brain was now lowest (ranked fifth) and skin was fourth.

### Effects of epigenetic reprogramming are module specific

One of the most remarkable recent observations from the epigenetic clock literature is the ‘resetting’ phenomenon via epigenetic reprogramming. For instance, numerous studies have demonstrated that reprogramming of adult somatic cells into iPSCs via Yamanaka factors can revert the epigenetic clock score back down to the age equivalence of fetal or embryonic cells ^12,23–26^. However, the degree of this reset is not perfectly consistent across clocks (Figure S5). Some clocks, like the original Pan-Tissue (Horvath clock), show very dramatic resetting, while others like DunedinPoAm or EpiTOC2, actually exhibit increases in epigenetic age. Additionally, there are some, like Hannum and GrimAge, that appear to increase during the initiation phase before changing their trajectory and then decreasing as cells mature into fully reprogrammed iPSCs. Thus, we tested whether these inconsistent results could be explained by module-specific differences—either the difference in the degree of signal from reprogramming sensitive modules and/or the differential weights of these modules.

We find two modules—green-yellow and green—that tend to be the most sensitive to resetting by epigenetic reprogramming. For six of the eight clocks shown in Figure 5, these two modules show marked decreases in epigenetic age, starting as early as the initiation period. The two clocks that do not show this are EpiTOC2, which does not contain either of these modules, and DunedinPoAm, for which the green-yellow module actually increases very dramatically. This result could be due to inverse weighting of the green-yellow module in the DunedinPoAm clock. Interestingly, DunedinPoAm and EpiTOC2 were also the clocks that exhibited contradictory results when examining reprogramming across the full clock scores. Another module that shows resetting in response to reprogramming is the light-blue module, which moderately decreases in all the clocks that it is present in, although to a lesser degree. The increase in EpiTOC2 is related to both pink and navy which make up most of EpiTOC2 but contribute very little to the other clocks. For the remaining modules, there seem to be less consistent and/or weaker patterns of change as cell convert from adult somatic cells (in this case, dermal fibroblasts) to iPSCs.

**Figure 5:**
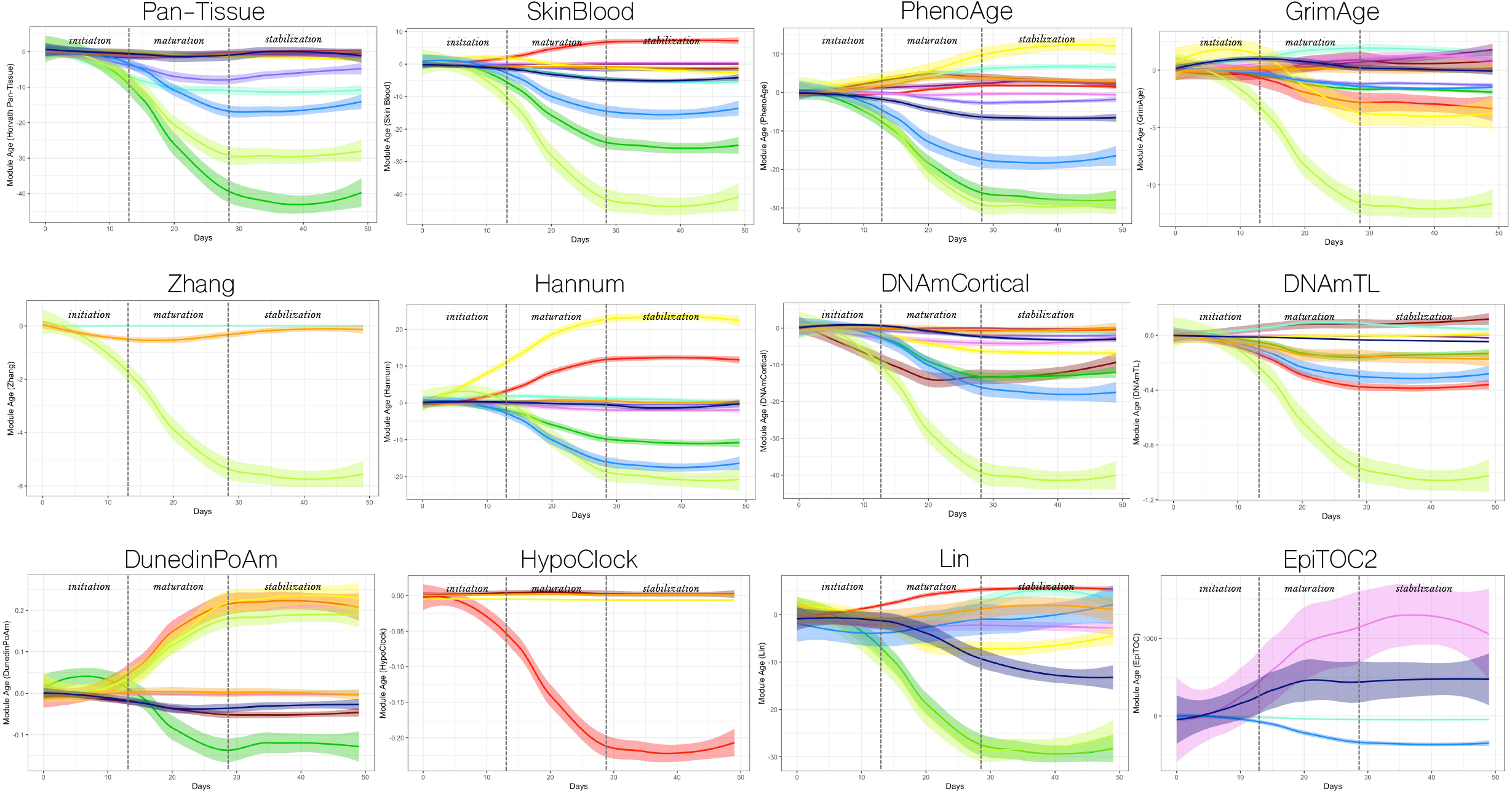
Clock module changes during time-course epigenetic reprogramming in fibroblasts (TRA-1-60 (+) cells) Module specific DNAmAge were calculated for eight clocks and plotted as a function of time (days) during reprogramming. Colors denote modules plotted for smoothed conditional means (loess) and shaded confidence intervals. Dashed vertical lines depict transitions between initiation, maturation, and stabilization phases. Y-axis depicts the degree of DNAmAge changes since baseline (day 0). We observe strong resetting in some (greenyellow, green, and light blue), but not all modules.

### Modules vary in predictions of mortality risk and associations with health related traits

When assessed in blood, epigenetic clocks have been shown to be predictive of remaining life expectancy. This is especially true for what have been referred to as 2^nd^ generation clocks, which were trained as predictors of aging outcomes (like mortality or clinical indicators)^13,19,22^ in contrast to 1^st^ generation clocks that were developed as chronological age predictors^12,29,31^. As with the other associations, we tested whether certain modules were capturing the life expectancy signals in epigenetic clocks. To do this, we used data from n=2,478 participants in the Framingham Heart Study (FHS), and adjusted for chronological age is a proportional hazard model of time-to-death. We find (Figure 6) that as with reprogramming, the green-yellow module is the most robust predictor of mortality risk when considering various epigenetic clocks overall (based on average z-score across all clocks). For instance, the green-yellow module has z-scores for mortality prediction greater than 8.8 for both the DNAmPhenoAge and the Zhang clock, followed by z-scores of approximately z=6.0 for GrimAge and DNAmCortical^35^. Also as with reprogramming, the largest outlier for the green-yellow module association with mortality is DunedinPoAm, which has a z-score of only z=1.0.

**Figure 6:**
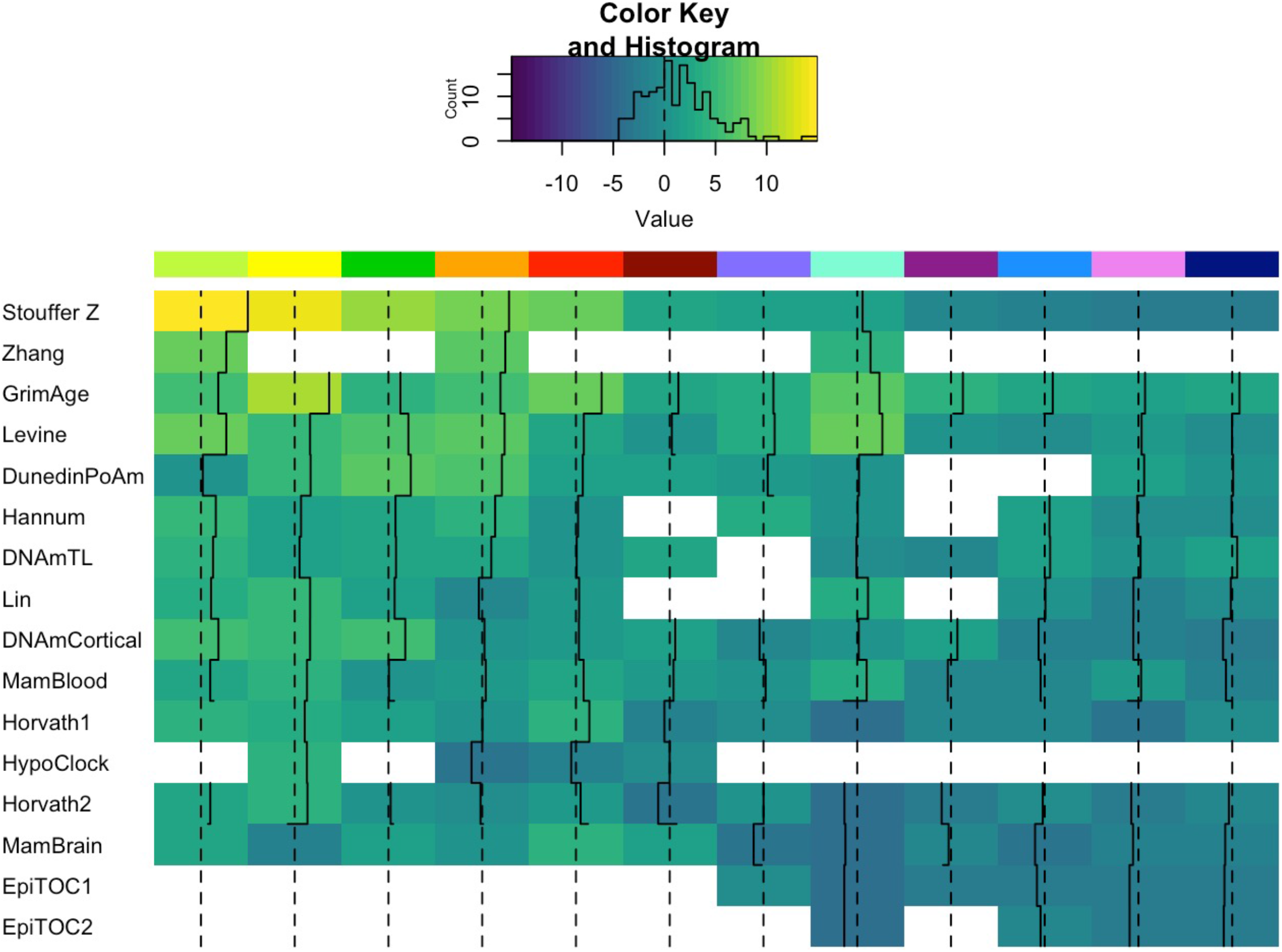
Mortality z-scores for epigenetic clock sub-modules. FHS whole blood data was used to test predictive associations between clock modules and allcause mortality. We plotted standardized z-scores for the effect of each clock module based on a cox proportional hazard model, adjusted for age and sex. The first row depicts the meta z-score estimated via Stouffer’s method. Modules (denoted by color on the top of each column) are arranged from highest meta-z (left) to lowest (right). Rows depict the epigenetic clock from which each module was estimated. Overall, the greenyellow module shows the strongest mortality prediction. However, the strongest individual module-clock pair was the yellow GrimAge module.

The highest overall mortality association for any single clock module was found for the yellow GrimAge module (z=11.7), although, it should be noted that GrimAge was actually trained in FHS data and is therefore overfit to some degree^19^. Interestingly, the z-score for the yellow module alone is only moderately weaker than the mortality z-score we observed for the full GrimAge clock (z=14.1, data not shown). Overall, this suggests that a large proportion of the mortality score is captured by this single module; although the other modules also contribute, with z-scores ranging from z=2.31 (navy) to z=8.7 (red). Like GrimAge, the DunedinPoAm clock has also been linked to mortality risk in the literature. When considering its module components, we find that the strongest mortality predictor is the green module at z=7.7, followed by the orange module (z=6.7). First generation clocks, like Hannum and Pan-Tissue, have been shown to predict mortality, albeit to a much weaker degree than second generation clocks^19,36^. Interestingly, both clocks have modules with strong mortality predictions. However, these seem to be counteracted by a number of modules with weak, or even inverse associations with mortality. For instance, the z-score for the green-yellow module is greater than 4 for both clocks—Hannum (z=5.0) and Pan-Tissue (z=4.7). However, four of the ten modules that make up Hannum have z-scores less than one (including one module with a negative z-score). This is even more dramatic for the Pan-Tissue clock, which has eight of twelve modules with z-scores less than 1, six of which are negative.

A few modules exhibit very weak, if not mostly inverse associations with mortality. These include the navy, pink, and light-blue modules. For the navy module, it shows a negative association with mortality in eight of the thirteen clocks that it is present in. The pink and lightblue modules exhibit negative mortality associations for six out of thirteen, and seven out of twelve clocks, respectively. Thus, the presence of these modules in epigenetic clocks could diminish their utility in estimating remaining life expectancy or mortality risk.

Using data from the FHS, we tested health associations for modules from three second generation clocks. Correlations between traits and clock modules (residualized for age and sex) are shown in Figure S6. For GrimAge, the most striking associations are with smoking, which are observed to be very strong for both the yellow and the red modules. We also find strong associations with cardiovascular related outcomes for greenyellow, cyan, yellow, red, green, and lilac (in descending order). Interestingly, the lilac is particularly associated with risk of coronary heart failure, but less associated with cardiovascular disease or coronary heart disease. The greenyellow module also appears to have the strongest association with age at menopause (suggesting higher epigenetic age associated with earlier menopause). For the PhenoAge modules, the green and greenyellow modules show the strongest smoking associations— although they are substantially lower than what was found for GrimAge modules. They also show the strongest associations with cardiovascular related outcomes and again, greenyellow is negatively associated with age at menopause. Finally, for DunedinPoAM, strong smoking associations are found for the orange module, followed by the green, yellow, and navy module. Cardiovascular related associations are found for red, orange, green, lilac, and navy. For this clock however, we find a reverse association for greenyellow consistent with what was observed in both the reprogramming experiment and for the mortality associations—suggesting that greenyellow signal is reverse coded in the DunedinPoAm clock.

### Specific Clock modules can distinguish tumor from normal tissue and track replication in vitro

We have previously reported that epigenetic age is accelerated in tumors compared to normal samples from the same tissue of origin—but only for certain clocks^28^. Therefore, we tested whether this could be due to differences between modules when it comes to distinguishing neoplastic vs. “normal” tissue. To test this we compared matched samples from five tissue types—breast, lung, pancreas, colon, and thyroid. Figure 7A shows the z-scores for each clock module regressed on cancer state (i.e. tumor vs. normal) and adjusted for age and tissue type. Points represent the different clocks and positive values suggest age acceleration in tumor vs. normal. Overall, we find that two modules— navy and light blue—show strong epigenetic age increases in tumor vs. normal samples. This was not driven by a single tissue type, as we found that it remained consistent even when stratifying by tissue (Fig S6). Next, we performed PCA on all navy clock module DNAmAge scores (e.g. Pan-Tissue navy, PhenoAge navy, etc.) and all light blue clock modules and then asked whether PCs could distinguish normal vs. tumor (Figure S7). For both modules, PC1—particularly for the navy module—can mostly segregate the two origin sample types, with only minimal overlap. When using PC1 for both modules, the distinction between tumor and normal is substantial, suggesting that across the five tissues, there is a consistent epigenetic aging effect as they transition to a solid tumor state. Thus, even though the overall epigenetic clock scores often cannot separate tumor vs. normal, when restricting to the parts of the scores captured by the navy and light-blue module, we can now discriminate tumorigenesis.

**Figure 7:**
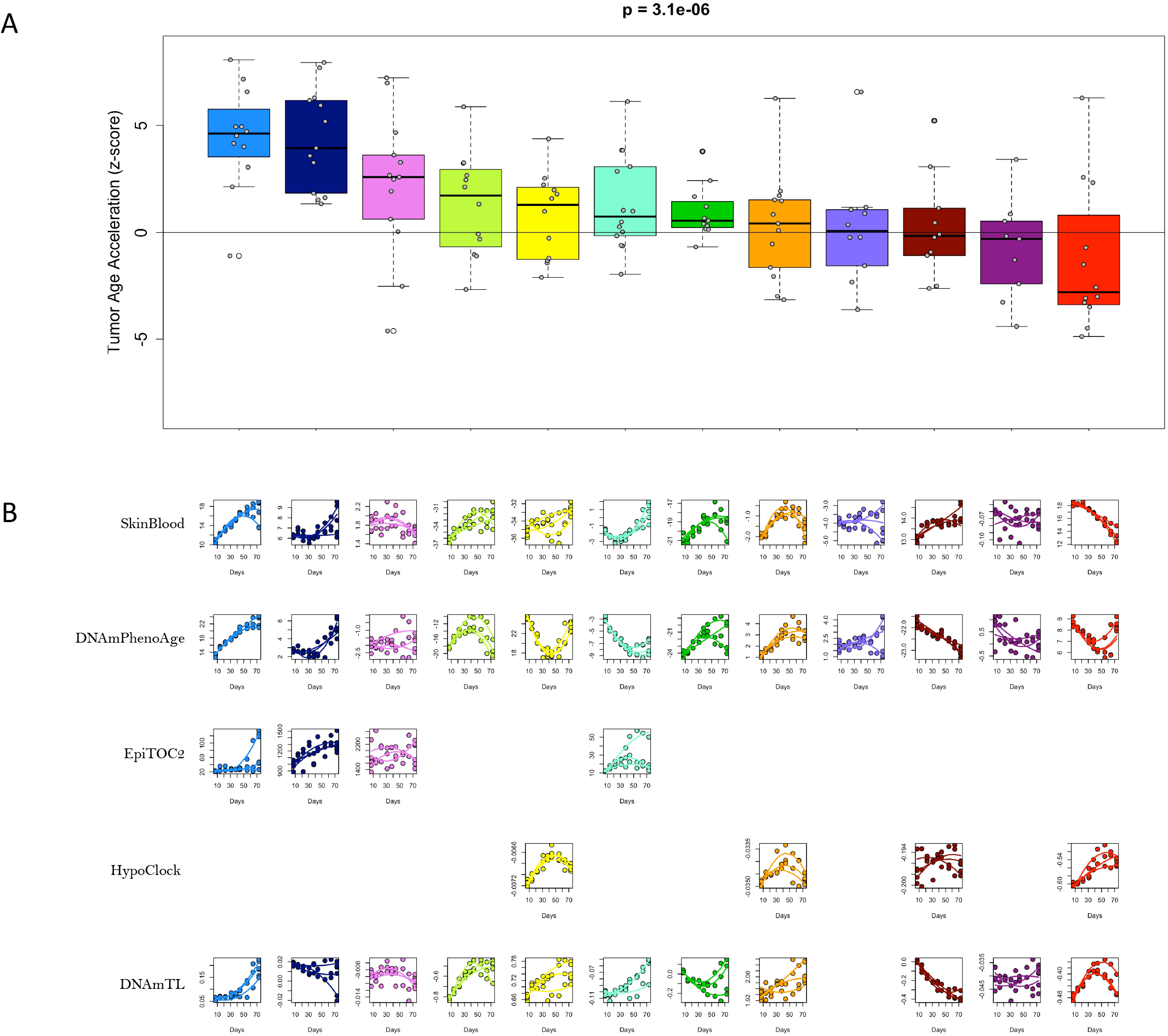
Tumorigenic clock module acceleration and associations with proliferation. (A) Data from clock modules estimated in tumors and normal tissues, for five tissue types (breast, colon, lung, pancreas, and thyroid) were used to test for accelerated DNAmAge in tumor. The z-scores of effect size (increase in DNAmAge for tumor versus normal tissue) for each clock module pair were extracted from a linear regression model, adjusted for age and tissue type. Boxplots, with means and standard errors are shown for z-scores pertaining to each module (denoted by color), with positive values suggesting accelerated aging in tumor, and negative values suggesting decelerated aging in tumor. P-value represents significant differences between modules using Kruskal-Wallis test. (B) Fetal astrocytes were passaged for 73 days and DNAm modules were estimated for three biological replicates at each passage. We plotted the change in module DNAmAge for five epigenetic clocks (rows are clocks and columns are modules, denoted by color). Second-degree polynomials were fit to infer change in module clock DNAmAge with passaging.

One explanation for this may be increase in proliferative capacity of tumors. Using a cultured fetal astrocyte model, we tested whether module-specific epigenetic age for five clocks (SkinBlood, DNAmPhenoAge, EpiTOC2, HypoClock (reverse coded), and DNAmTL (reverse coded)) tracked with days in culture. Results show that, across all four clocks containing the light blue module, the score increases over the course of the passaging experiment (Figure 7B). The navy module also shows increases with passaging for all but the DNAmTL clock. Other interesting trends include the increase in the green-yellow module, which hits an inflection point around 43 days in culture (passage 6). This is noteworthy, as this was the point in time when we began to observe an up-tick in senescence markers and slowing of cell population doublings (cPD), suggesting cells are beginning to enter a state of growth arrest (Figure S8). The orange module also appears to display this trend, particularly in the SkinBlood and DNAmPhenoAge clock.

As mentioned previously, the two mitotic clocks—EpiTOC2 and HypoClock—were developed to track these types of changes and generally exhibit increases in module scores as a function of passaging. In the EpiTOC2 clock, the navy module shows the greatest degree of acceleration, while passaging-related changes are also seen in the light-blue module (as mentioned above), but not the pink module. For HypoClock, red and yellow show substantial increases, while changes are less robust for orange and dark-red. The yellow, orange, and dark-red modules of the HypoClock also tend to revert around passage 6, likely due to growth arrest in senescent cells. Finally, the telomere length clock (DNAmTL) shows increases in the light-blue, green-yellow, yellow, cyan, orange, and red modules. The dark-red module also shows strong change, but in the incorrect direction (indicative of lengthening of telomeres as a function of passaging).

### Shared module signals

Using the module clocks measured in whole blood, liver, adipose, skin, and brain, we performed consensus clustering to test whether clock module pairs were picking up shared signals. Affinity matrices based on correlations were calculated and were combined such that the minimum (greatest distance) was selected for the consensus matrix. We then performed hierarchical clustering on the consensus matrix (Figure 8). We identified two clusters of clock modules, which show distinct association trends. For instance, the first cluster included light-blue, navy, pink, and some cyan modules. The EpiTOC clocks, DNAmCortical, and mammalian clocks were observed to group tightly, while the Pan-Tissue, SkinBlood, Levine, Lin, and GrimAge clocks tended to group in this cluster. In general module clocks in this cluster exhibit non-significant or weakly negative associations with mortality risk, strong acceleration in tumor versus normal tissue, and moderate rejuvenation during reprogramming—mostly restricted to the maturation phase. The other cluster was observed in the greenyellow, green, yellow, and some cyan modules. This cluster of module clocks tended to show strong prediction of mortality risk, moderate acceleration in tumor versus normal tissue, and stronger resetting in reprogramming— mostly during maturation phase.

**Figure 8:**
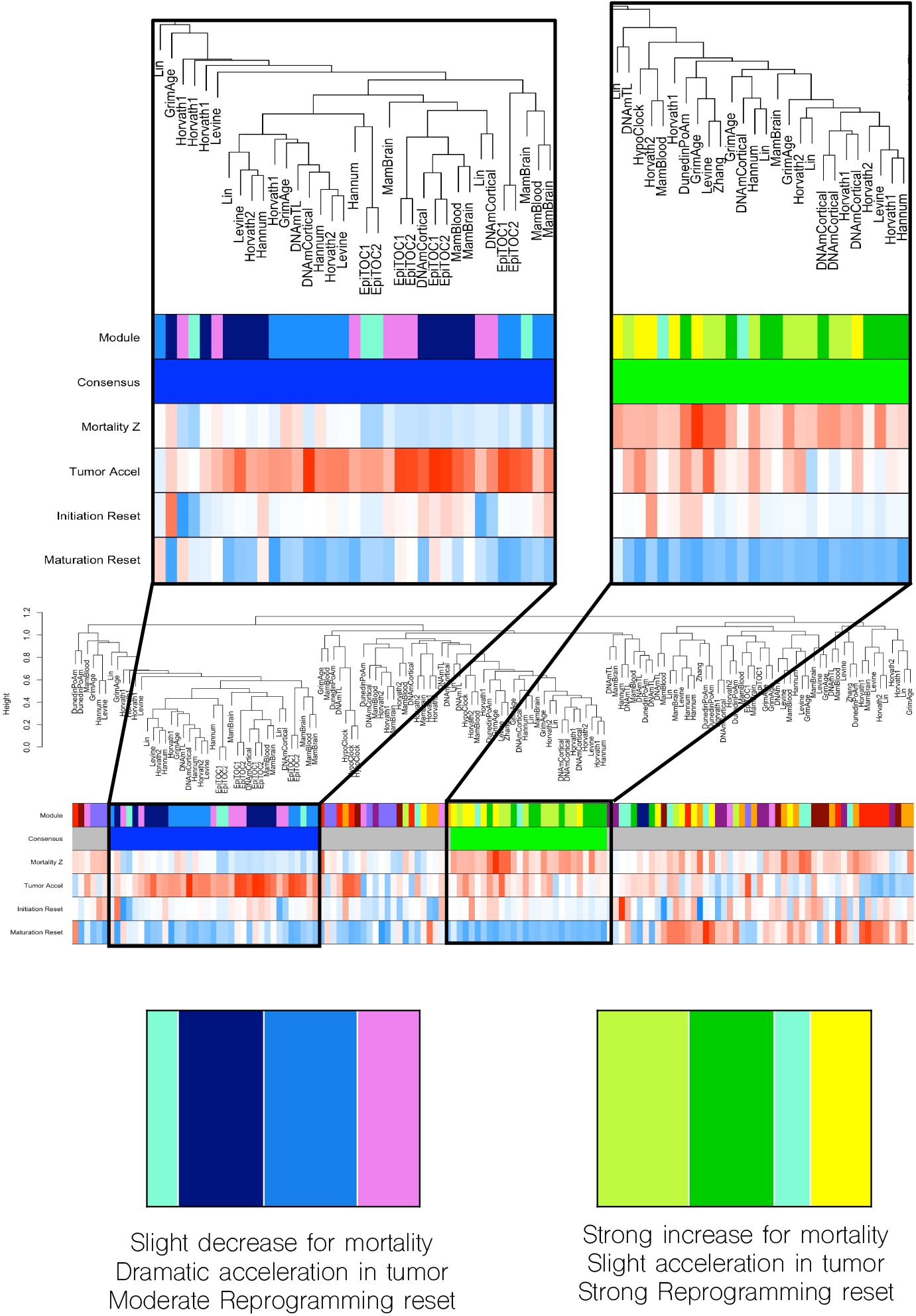
Tumorigenic clock module acceleration and associations with proliferation. Consensus clustering of clock modules was performed using data from whole blood, liver, adipose, skin, and brain. We identified two group of clock modules (enlarged on the top row), which show distinct association trends. For instance, the light-blue, navy, pink, and some cyan modules tend to cluster and exhibit non-significant or weakly negative associations with mortality risk, strong acceleration in tumor versus normal tissue, and moderate rejuvenation during reprogramming. Conversely, the green-yellow, green, yellow, and some cyan modules tended to cluster, show strong prediction of mortality, moderate acceleration in tumor, and strong resetting in reprogramming. All remaining module clocks are shown in the middle dendrogram. On the bottom row, the relative sizes of the bands for each color reflect the degree of representation in that cluster. For instance, cluster one is primarily the light-blue and navy, with some inclusion of pink and cyan.

## DISCUSSION

Since their emergence in 2011^10^, epigenetic clocks have been feverously applied to study a wide range of scientific questions related to aging and longevity, cancer, infectious disease, development, and social and environmental determinants of health^27^. Yet despite their mass-appeal, our understanding of what makes them tick is rudimentary. Very little is known when it comes to the pathways or mechanisms involved in the age-related changes in DNAm, and even less is known about why biomarkers built from these changes are associated with risk for morbidity/mortality. Different clocks often do not agree after regressing out chronological age, and the underlying logic that explains these discrepancies remains unknown. Some studies have begun to use in vitro models to try to uncover the underpinnings of epigenetic clocks, but so far a lack of consensus remains^37^. Reflecting epigenetics hypothesized position as a master conductor of cellular traits, it follows that clocks derived from DNAm will integrate multiple distinct mechanisms of age-related change, rather than proxying a singular distinct mechanism. Different mechanisms thus influence different the various DNAmAge scores and it is only after deconstructing clocks to identify their respective “pieces” that we can start to understand the inner workings of epigenetic aging.

Clock CpGs were clustered into twelve distinct modules. Overall, these modules exhibited differences in their dynamic ranges with aging in multiple tissues and biofluids and were differentially responsive to epigenetic reprogramming of fibroblasts. They also showed differing enrichment in their proximity to high versus low CpG density regions. For instance, the greenyellow module—which as we will discuss, is one of the most interesting clock modules—tended to be in low CpG density, intergenic, Open Sea regions. CpGs in this module did not tend to show extreme hyper- or hypomethylation when measured in tissues or blood, but instead tended to have dynamic ranges closer to 0.5, suggesting greater heterogeneity between cells. The one exception was in reprogrammed fibroblasts, for which CpGs in the green-yellow module tended towards hypermethylation. Furthermore, despite having DNAm levels around or just under 0.5, this module showed large decreases in DNAm as a function of age. For instance, mean DNAm for green-yellow CpGs was 39% in blood from 20-39 year-olds, 36.6% in blood from 50-59 year-olds, and 34.4% in blood from 70+ year-olds (p=0.002, data not shown).

If DNAm changes were purely reflecting entropic alterations or epigenetic drift, we would expect to see a bias against changes in CpGs that start around 0.5 (corresponding to random chance of methylation at a given site)^38^. However, what we observe is actually a regression away from the mean, in which these heterogeneous populations of cells are systematically losing DNAm with time. This suggests that the green-yellow module’s notable pattern of epigenetic aging is unlikely to stem from noise or aberrant DNAm changes with age. Instead, DNAm changes may reflect cellular selection pressure or clonal expansion in which the cells without DNAm at these CpGs are able to outcompete (proliferate more than) the ones with DNAm^39^. Alternatively, it could reflect a regulated compensatory mechanism that gets initiated with aging,^3^ or a continuation of a developmental program that is not turned-off^40^. These scenarios have different implications for our understanding of epigenetic changes. The first would suggest that individual cells are not changing DNAm patterns with age, but rather the changes that are observed in bulk data are happening at the level of cell populations, shifting prevalence of cells with heterogenous states. The second and third scenarios, on the other hand, would suggest within cell DNAm changes, perhaps as a response to extracellular environment or signaling changes with aging (e.g. integrated stress response (ISR)^41^), or as an extended developmental program that fails to be extinguished—somewhat aligned with the hyper-function theory of aging^40^. In moving forward, single-cell DNAm data may help distinguish individual vs. population changes.

While the green-yellow module does not support an epigenetic noise hypothesis, there were modules for which the patterns of DNAm changes could be indicative of drift. For instance, the red and yellow modules were hypermethylated in iPSCs and young tissues and showed decreases in DNAm with age. Conversely, the light-blue module was hypomethylated in iPSCs and young samples and tended to gain DNAm with aging. These both exhibit regressions towards the mean, suggestive of increasing cellular heterogeneity and stochastic gains or losses of DNAm with aging. Thus, DNA methylation aging could be consistent with multiple aging theories that manifest in distinct CpG modules.

Given the systematic differences in behaviors—and likely mechanisms—between modules, we hypothesized that this could account for differences observed between clocks. Using the published clock equations, we were able to estimate clock scores that only captured the part of the overall signal due to each module. Interestingly, we found that clocks varied in the degree of influence each module contributed towards the overall clock score. For instance, first generation clocks, like Pan-Tissue and SkinBlood, which were trained to predict chronological age in diverse sample types had very similar module distributions. This was distinct from two second-generation clocks—GrimAge and DunedinPoAm—which were trained in whole blood to predict mortality^19^ or physiological changes with aging^22^. The one exception to this pattern was DNAmPhenoAge, which despite being considered a second-generation clock, comprised module distributions that were more similar to first-generation clocks.

This is potentially noteworthy given that DNAmPhenoAge was trained to predict an aging measure derived from clinical chemistry variables (like DunedinPoAm) and that was developed to predict remaining life expectancy (like GrimAge). However, unlike GrimAge and DunedinPoAm, the dependent variable (or ‘ground truth’) used for training DNAmPhenoAge, also included chronological age^13^. DunedinPoAm, on the other hand, was trained in a cohort of same-aged individuals^22^, while GrimAge included observed age in its derivation on the other side of the equation—as a predictor along with DNAm—essentially adjusting out its effects^19^. This is consistent with the modules’ age and mortality associations: As GrimAge and DunedinPoAm do not predict chronological age from methylation, they utilize less information from navy and light blue (age-associated but weakly mortality-associated), but more information from red and orange (mortality-associated but weakly age-associated). On the other hand, DNAmPhenoAge utilizes all these modules. It has also been shown that like first generation clocks, DNAmPhenoAge, performs well when assessing perturbations in cultured cell models^28^, which may insinuate that it is more of a hybrid between first and second generation clocks.

Three other clocks also exhibited interesting distributions of modules—such that they were almost entirely captured by just two modules. For instance, Zhang^42^, which is a mortality clock (but without age adjustment), derived approximately 80% of its signal from just the orange and green-yellow modules which are both mortality-associated across many clocks. HypoClock, which was developed to capture mitotic history based on increasing hypomethylation at solo-WCGW sites was made up almost entirely of just the red and dark-red modules; while another mitotic clock, EpiTOC2—developed from promoter associated CpGs in islands marked by the polycomb repressive complex-2 (PRC2)^34^—was predominantly derived from the navy and pink modules.

Given that different clocks are composed of differing distributions of modules, we hypothesized this may explain the discordant results often observed when utilizing multiple distinct clocks simultaneously to try to answer the same scientific question^21,27,28,43^. This has been a major roadblock in the epigenetic clock science given that there is no ground truth for the latent concept of ‘biological age’ that each clock is attempting to proxy. As such, discordant association results have weakened the field’s perceptions of the validity of epigenetic clocks. For instance, if one were to run a clinical trial and assess efficacy using epigenetic clocks, it would be difficult to make a claim about intervention effects if different clocks did not agree on whether epigenetic age was decreased, increased, or unchanged. However, this could happen if an intervention targets only a specific domain of epigenetic aging and that domain is more or less captured by different clocks.

To test this possibility, we assessed the effect of a popular emerging aging intervention— reprogramming—on module specific epigenetic age scores. We found that only specific modules were reset in response to reprogramming, while others remained unchanged, or even increased. For instance, the green-yellow module showed one of the greatest “reset” responses to reprogramming. The one exception to this was the DunedinPoAm clock. In this clock, the greenyellow module still showed a large effect but in the opposite direction, indicative of increased epigenetic age. This is likely due to opposite weighting of these CpGs in the DunedinPoAm clock compared to other clocks (e.g. increased DNAm in green-yellow module CpGs is reflective of decreased age scores in most clocks, but increased age scores in DunedinPoAm). It is unclear why green-yellow CpGs in DunedinPoAm are reverse weighted, but one explanation may be that training was conducted in age invariatnt samples. Importantly, the decline in epigenetic age for the green-yellow module began during the “initiation” phase, suggesting that some degree of epigenetic age resetting may precede the dedifferentiation that occurs over the course of the maturation phase^23^.

Other modules that showed consistent decreases in epigenetic aging in response to reprogramming were the green module and the light blue module. Many, however, showed little change, or on occasion, increases. This is critical to consider when inferring the potential implications of any interventions. Behavioral and therapeutic interventions that have used epigenetic clocks have repeatedly made claims about slowing or reversing biological age in response to decreases in epigenetic clock scores. However, epigenetic clocks are not perfectly synonymous with biological age and problems could arise if modules with no bearing on lifespan and healthspan are the only ones responding to a given intervention. Most epigenetic clocks, especially second-generation clocks, are predictive of a variety of aging outcomes, but that does not mean these associations are robust across all CpGs in the clock. In recent studies, when a given intervention has been shown to alter clock scores, there have been assumptions made that the intervention will in-turn influence outcomes like morbidity and mortality given that the clock is predictive of them. However, if clocks are made-up of distinct modules—modules A, B, and C for purposes of this example—one could imagine a scenario in which module B is responsive to an intervention and yet it is module A that is predictive of outcome. In this example, the intervention likely has no bearing on the outcome of interest.

Fortunately, when it comes to the two modules that are most responsive to reprogramming, they do show strong predictions for mortality risk. The green-yellow module was the strongest predictor of future mortality risk (on average across different clocks), while the green module also had strong effects. Unlike the scenario above, the concept that an intervention will impact outcomes because it impacted clock scores has more grounding, given that all CpGs within a module are multi-collinear and thus likely to change synergistically. However, there were also modules that did not typically change with reprogramming but were shown to be predictive of mortality; one example being the yellow module. Yet, the strongest association between mortality risk and a yellow module was found for GrimAge. This clock was also the singular case in which the yellow module exhibited clear age reversal, although not beginning until the maturation period. However, in both DNAmPhenoAge and DunedinPoAm, the yellow module was increased with reprogramming, but is inversely associated with lifespan, suggesting that reprogramming contributes to a “higher risk” profile for this module. In this case, it is not clear what the change in mortality risk could be inferred for interventions that had a beneficial effect on some modules (e.g. green-yellow and green), but neutral or deleterious on others (e.g. yellow). Regardless, although the clocks have been associated with mortality risk and are responsive to reprogramming, it has yet to be proven whether changes in a given clock (or module) will garner a change in mortality risk. For instance, if clocks are read-outs of other stressors, rather than causes of aging per se, and if these stressors are orthogonal to the mechanisms being targeted by reprogramming, then it is unlikely that reprogramming would restore a system to a state with greater resilience and less risk. That being said, results from in vivo reprogramming studies do suggest that the reversal of the epigenetic aging profile may at least be accompanied by improved health. While more direct evidence of the role of specific DNAm changes is needed, the modules may speed up identification of critical epigenetic mechanisms, by highlighting CpGs for which attention should be focused.

Another important, yet often overlooked application of epigenetic clocks, are the tissue-asynchronous changes over time. For instance, the original Horvath Pan-Tissue clock was developed to estimate age accurately across nearly all tissues^12^. However, it makes sense that tissues and cells would not all age at the same rate. Interestingly, we found that when we consider the modules for this clock, rather than the clock as a whole, tissue differences are suddenly apparent. For a given tissue, age is overestimated by some modules, but underestimated by others. Thus, when modules are summed to derive the full clock score, these differences average out and tissue aging patterns appear homogenous. It is interesting to note that when considering the different modules, they are not unanimous in estimating which tissues are older versus younger. For instance, the Pan-Tissue green-yellow module puts adipose as the oldest and brain as the youngest, while the pink module ranks brain as oldest and skin as youngest. In moving forward, it will be essential to determine what tissue or cell specific aging factors produce these differences, or if this is simply an artifact of the training method. For instance, tissue residency^44^, metabolic rates^45^, oxygenation^46^, or proliferation rates^47^ could contribute.

We also observed that modules differed in their trajectories of change during growth and development. Many of the well-studied epigenetic clocks have been shown to exhibit exponential increases as a function of age, when measured in fetal or pediatric samples^48^. However, this is not the case for all modules. Those like the green and green-yellow module do show very accelerated changes during this developmental period, while the navy and pink modules exhibit more linear change, albeit with high variance in fetal samples. This was also a puzzling finding when examining two clocks (EpiTOC2 and HypoClock) that were intentionally developed to track mitotic rates. None of the modules in either clock exhibited drastically accelerated developmental change, possibly because they were trained in adult blood.

Furthermore, their ordering of tissue asynchronous aging rates were not always in line with what has been reported in regards to tissue differences in turnover. One might expect that mitotic clocks would consistently rank brain as the “youngest”, given that many brain cells—neurons and to some degree astrocytes—are generally postmitotic in adults^49^. However, this was only observed in one of the three modules for each clock (light-blue for EpiTOC2 and orange for HypoClock). Moreover, these specific modules were found to only contribute tiny fractions to the overall clock scores (2% and 4%, respectively).

Taken together, this may suggest that EpiTOC2 and HypoClock are not capturing proliferation per se, but something else correlative with turnover within a given tissue or cell type. Further evidence of this is that the modules in these two clocks did track with passaging in astrocytes. For EpiTOC2 the navy, light-blue, and cyan modules, but not the pink module, showed strong increases as a function of passaging, or days in culture. Interestingly, the light-blue and navy modules exhibited an exponential increase across the lifespan of these cells. This suggests they are tracking phenomena like genomic instability or activation of senescence pathways. In contrast, other modules displayed a u-shaped trajectory (green-yellow, yellow, green, orange, red, and dark red). This included most HypoClock modules. For these, we observed an inflection point that tended to occur around the time when cPD for these cell lines decelerated and the percent β-galactosidase positive cells increased. One possible explanation for this sudden deceleration is that it may reflect heterogenous selection pressures or fitness, in which less resilient cells undergo apoptosis or senescence, while more resilient cells remain proliferative in the population for longer and thus contribute more to the average signals at later time-points. However, single-cell analyses will be needed to determine whether these different trajectories reflect cell intrinsic changes or population shifts.

The modules that increased most dramatically with passaging were also the most accelerated in solid tumors in contrast to normal tissue. This was consistent across multiple tissue/tumor types, including breast, pancreas, colon, thyroid, and lung. The trend was also mostly clock independent, since we observed a consistent trend across all navy sub-clock modules and the same for all but one (Lin) of the light-blue sub-clock modules. It remains unclear whether these changes develop after tumor initiation because of driver mutations or increased proliferation, or instead whether normal tissues that have acquired them become more tumorigenic and thus prone to transformation. The latter would suggest that such aging-related epigenetic changes may promote oncogenicity and could be one additional explanation for the exponential increase in cancer incidence with age^50^. Despite their association with cancer, these modules did not show associations with mortality when measured in blood using the FHS study. This may be because acceleration will only promote tumorigenesis in the tissue for which it is observed and thus acceleration in blood would not be indicative of acceleration in tissues and organs for which more lethal solid tumors arise.

Overall, this study is the first to show that by deconstructing epigenetic clocks into functional modules, we were able to compare differences more directly between the various clocks that have been developed. Differential proportions of module signals in clocks contribute to clock disagreement, which would account for some of their inconsistencies regarding associations with outcomes or mechanisms. We also identified potentially promising clock modules, such as the green-yellow module, which shows dramatic resetting in response to reprogramming, is strongly associated with mortality risk, exhibits tissue asynchronous aging trends that may reflect proliferation rates, and is altered with in vitro passaging of cells. This module was found to be enriched for intergenic CpGs that showed non-entropic change with age, suggesting that it may reflect the maladaptive consequences of programmatic alterations or clonal expansions. Moving forward, it will be critical to determine the exact mechanistic drivers of these types of epigenetic age changes and better understand the regulatory features that account for the documented associations with risk of morbidity and mortality.

## METHODS

### PhenoAge, GrimAge and Mammalian clock training

Multiple semi-novel epigenetic clocks were generated for this study. The first was a novel version of DNAmPhenoAge, motivated by our observation that the original DNAmPhenoAge is a noisy measure among epigenetic clocks^51^. We found that simply increasing the training sample size was sufficient to increase reliability to the level of other clocks. Therefore we combined data from the InCHIANTI and HRS datasets to train a novel 960-CpG DNAmPhenoAge clock to predict clinical PhenoAge from DNA methylation data^13,52^. Note that this CpG-based DNAmPhenoAge is distinct from the highly reliable principal component-based PCPhenoAge reported recently, though both were trained using the same samples and same 78,464 CpGs as input. We also trained a surrogate estimate of the GrimAge clock. CpG identities and weights for the original GrimAge clock are not published and thus we recreated them in order to have these parameters^19^. To do this we calculated GrimAge and all of its components in the Framingham Heart Study using the online epigenetic clock calculator (http://dnamage.genetics.ucla.edu/new). We then predicted each of the components from chronological age, sex, and the 30,084 CpGs used as input to the online calculator. This was carried out utilizing elastic net regression with 10-fold cross validation and an elastic net mixing parameter alpha of 0.5, utilizing the glmnet R package. We then predicted the full GrimAge score from chronological age, sex, and the individual GrimAge components, which closely agreed with the published weights of the GrimAge components in the final calculation. 944 total CpGs were included in this version of GrimAge, similar to the originals’ 1,030 CpGs.

To facilitate application of our modules to future mechanistic studies in animal models, we included age-associated CpGs highly conserved across mammalian species^53^. To do this, we selected 1,990 CpGs present on the human 450K array, EPIC array, and the new Mammalian Methylation Array. We identified CpGs that correlate with age with absolute biweight midcorrelation greater than 0.2 in whole blood (GSE40279; 214 CpGs), brain (GSE74193; 227 CpGs), liver (GSE61258 selecting non-smoking and non-diseased samples; 392 CpGs). We also identified CpGs that correlate with age with absolute biweight midcorrelation greater than 0.15 in skin (GSE90124; 113 CpGs) and adipose (MT1866; 202 CpGs). The majority of these CpGs were tissue-specific, resulting in a total list of 820 CpGs. We also trained mammalian clocks using all 1,990 CpGs as input to predict age utilizing elastic net regression with 10-fold cross validation and an elastic net mixing parameter alpha of 0.5, utilizing the glmnet R package. We trained such a 293-CpG mammalian blood clock in GSE40279 which we validated in the FHS cohort. We also trained a 149-CpG mammalian brain clock in GSE74193 which we validated in the Religious Orders Study. The majority of these CpGs overlapped with the age-correlated CpG sets, resulting in a total of 1,022 CpGs that were included in clustering and module identification.

### Module Identification

Seven DNAm array datasets were used to group clock CpGs into modules. These included DNAm data from whole blood (FHS cohort), liver (GSE48325), brain (GSE74193), skin (GSE90124), adipose (MT1866), reprogrammed fibroblasts (GSE54848), and cultured fetal astrocytes (primary data). For each of the datasets, we restricted CpGs to n=5,717 that have been included in existing epigenetic clocks. We then estimated a signed adjacency matrix based on biweight midcorrelations between CpG pairs^54^. Additional, we also estimated adjacency based on Euclidean distance for two of the datasets—whole blood, and fibroblast reprogramming. For this step, we did not standardize values given that CpGs all utilize the same unit of measurement (DNAm beta values), which holds biological meaning, suggestive of percent DNAm. Adjacency matrixes were then converted to affinity matrices via nearest neighbors (n=500). Values were then standardized across affinity matrices to make datasets comparable, and finally combined into a Meta-Affinity matrix by taking the sample quantile at probability p=0.25 across matrices for each *p×p* cell. Our reasoning for including affinity matrices from correlation distances for all seven datasets, but using only two Euclidean based affinity matrices was to over emphasize the influence of strong correlations versus just having similar beta-values. This is because we are interested in dynamic changes that track together across perturbations—natural aging, reprogramming, and passaging. Spectral clustering was then carried out in accordance with traditional methods, by first, computing the unnormalized graph Laplacian (U=D–A), computing eigenvectors, and then performing k-means clustering. Given that our goal was to identify large clusters of strongly connected CpGs, we used a cluster size of k=600 for clustering and the pruned clusters with less than 100 CpGs, denoting them as unassigned. This resulted in the 12 large CpG modules.

### Module sub-clock estimates

In each of our datasets, we estimated the module-specific clock values for existing epigenetic clocks. To do this, we used the published corresponding weights for each CpG, but instead of multiplying by the weights and summing across all CpGs in a clock, we only summed across the CpGs in a given module. For instance, the generate a score for the Horvath Pan-Tissue red module, we took only CpGs in the Pan-Tissue clock that had been assigned to the red module, multiplied by their respective weights, and then summed across them. Thus, adding up all modules (in addition to weighted unassigned CpGs and the reported intercept for a clock) would produce the full clock score.

To estimate the proportion of signal captured by each clock, we summed the absolute value of all modules in a clock and then took the ratio of each module to the sum. The reason for taking absolute value is that some modules are actually negative, so they may have a large influence on the clock, but the variance is in the degree of deceleration for that module. The signal proportions were estimated in both whole blood and in fibroblasts during reprogramming given that CpG values multiplied by weights may change depending on data modality.

### Association testing

To test for CpG specific aging, passaging, and reprogramming effects, we estimated the degree of DNAm change as a function of one year change in chronological age, or one day change during passaging, or over the maturation phase of reprogramming (days 7-28). This was done by regressing DNAm for each individual CpG on the time variable (either chorological age in years or day of experiment). The beta coefficients were then used to indicate degree of temporal change. Aging changes were assessed in whole blood, adipose, liver, skin, and brain. For brain, samples were stratified by age such that effects were calculated for fetal samples (age<0) and adult samples (age>20) separately.

To test for temporal changes in clock, or clock modules, function of reprogramming in dermal fibroblasts, we first centered values so that the mean at initiation (day 0) would equal zero. Thus, changes in epigenetic age can be interpreted as change (increase or decrease) from baseline. We plot smoothed conditional means, with loess smoothing function, and shaded confidence intervals. Data was available for ten time points—days 0, 3, 7, 11, 15, 20, 28, 35, 42, and 49. Samples included sorted TRA-1-60 (+) cells after day 3. At day 28, cells were concerned fully reprogrammed into iPSCs and were passage three subsequent times in one week intervals.

FHS data from the offspring cohort was used to test for mortality association for each clock module. Data included n=2,478 sample, for which n=315 deaths were observed over 19,392 person-years. Cox proportional hazard models were used to test for prediction, with time-since baseline as the time-to-event variable, and non-deaths coded as censored at last observation. Models also included adjustment for chronological age. The z-score for each model was used to assess and compare the mortality associations between clock modules.

To test for association between modules and cancer state, we used ordinary least squares regression, controlling for chronological age and tissue type. Data included samples from breast (9 cancer, 4 normal); colon (12 cancer, 5 normal); lung (7 cancer, 3 normal); pancreas (6 cancer, 4 normal); and thyroid (70 cancer, 12 normal). Given that we had unequal distributions of tissue types, tests were also conducted in tissue-stratified models as a sensitivity analysis. As with mortality, z-scores were utilized to assess effect sizes given that clock (and especially clock modules) have different variances). A Kruskal-Wallis test was used to compare tumor age acceleration between different modules using clock-specific z-scores from the OLS model.

### In vitro passaging of fetal astrocytes

Finally, we also assessed changes in modules as a function of passaging in fetal astrocytes. A separate publication is in preparation concerning the astrocyte experiments. Briefly, three cell lines of fetal astrocytes were derived from cerebral cortex of the same donor (ScienCell #1800). Tissue was received by ScienCell Research Laboratories from non-profit tissue providers, obtained with informed consent of donor’s family aged over eighteen, and under established protocols in compliance with an institutional review board and local, state, and federal laws. No payment, commercial rights or financial rights were provided to the donor family. Further details can be obtained from ScienCell Research Laboratories.

Cells were exhaustively passaged and split a total of 10 times (9-15 cumulative population doublings, depending on replicate), where ß-gal activity (C12FDG) was measured using confocal microscopy at each passage to confirm exhaustive replication was achieved (data not shown). Cells were seeded at 8,000 cells/cm2 with appropriate growth media and supplements (complete astrocyte medium, containing amino acids, vitamins, hormones, trace minerals, 2% fetal bovine serum and 1% PEN/STREP in HEPES pH 7.4 bicarbonate buffer, ScienCell #1801) to promote cell adhesion and growth. Of note, Poly-L-Lysine was not required for adequate cell adhesion. Cells were grown under normoxic conditions (20% O2, 5% CO2) at 37°C.

Cells were split when they reached approximately 90% confluence or when static growth was achieved. Cells were counted using the Invitrogen countess and cell counting chamber slide with trypan blue. Cumulative population doubling was calculated using the initial and final cell density, as determined by the countess (2x=FD/ID, where x=population doubling, FD=final cell density and ID=initial cell density). Longitudinal samples were collected at every passage and DNA was extracted using the Qiagen DNAeasy Blood and Tissue Kit (69504). Note, samples were treated with proteinase K and RNAse A and eluted with 200 μl elution buffer. Following final elution, DNA was verified using nanodrop (Thermo Scientific). Spin concentration was used as necessary with low DNA content samples. Prior to library preparation we used a Qubit fluorometer (Thermo Scientific) to quantify the extracted genomic DNA. All samples were assigned a single-blinded code and randomized for library preparation and sequencing to control for any batch errors. DNAm data was generated using the Infinium HumanMethylation450 BeadChip and preprocessed using minfi45 and normalized using the noob method56. Scatterplots with fitted polynomials (to power 2) were plotted to visualize changes in clock modules over the experimental period.

## Notes

### Competing Interest Statement

The authors have declared no competing interest.

### Summary of Updates

Supplementary tables were previously not included

## CITATIONS

1 Kennedy, B. K. et al. Geroscience: linking aging to chronic disease. Cell 159, 709–713 (2014).

2 Novikoff, A. B. The Concept of Integrative Levels and Biology. Science 101, 209–215, doi:10.1126/science.101.2618.209 (1945).

3 Ferrucci, L., Levine, M. E., Kuo, P. L. & Simonsick, E. M. Time and the Metrics of Aging. Circ Res 123, 740–744, doi:10.1161/CIRCRESAHA.118.312816 (2018).

4 Lopez-Otin, C., Blasco, M. A., Partridge, L., Serrano, M. & Kroemer, G. The hallmarks of aging. Cell 153, 1194–1217, doi:10.1016/j.cell.2013.05.039 (2013).

5 Stueve, T. R., Marconett, C. N., Zhou, B., Borok, Z. & Laird-Offringa, I. A. The importance of detailed epigenomic profiling of different cell types within organs. Epigenomics 8, 817–829, doi:10.2217/epi-2016-0005 (2016).

6 Ahuja, N., Li, Q., Mohan, A. L., Baylin, S. B. & Issa, J. P. Aging and DNA methylation in colorectal mucosa and cancer. Cancer Res 58, 5489–5494 (1998).

7 Lu, Y. et al. Reprogramming to recover youthful epigenetic information and restore vision. Nature 588, 124–129, doi:10.1038/s41586-020-2975-4 (2020).

8 Jabbari, K. & Bernardi, G. Cytosine methylation and CpG, TpG (CpA) and TpA frequencies. Gene 333, 143–149, doi:10.1016/j.gene.2004.02.043 (2004).

9 Derreumaux, S., Chaoui, M., Tevanian, G. & Fermandjian, S. Impact of CpG methylation on structure, dynamics and solvation of cAMP DNA responsive element. Nucleic acids research 29, 2314–2326, doi:10.1093/nar/29.11.2314 (2001).

10 Bocklandt, S. et al. Epigenetic predictor of age. PLoS One. 6, doi:10.1371/journal.pone.0014821 (2011).

11 Horvath, S. & Raj, K. DNA methylation-based biomarkers and the epigenetic clock theory of ageing. Nat Rev Genet 19, 371–384, doi:10.1038/s41576-018-0004-3 (2018).

12 Horvath, S. DNA methylation age of human tissues and cell types. Genome Biol 14, doi:DOI: 10.1186/10.1186/gb-2013-14-10-r115 (2013).

13 Levine, M. E. et al. An epigenetic biomarker of aging for lifespan and healthspan. Aging (Albany NY) 10, 573–591, doi:10.18632/aging.101414 (2018).

14 Horvath, S. et al. Obesity accelerates epigenetic aging of human liver. Proc Natl Acad Sci U S A 111, 15538–15543, doi:10.1073/pnas.1412759111 (2014).

15 Horvath, S. & Levine, A. J. HIV-1 infection accelerates age according to the epigenetic clock. J Infect Dis, doi:10.1093/infdis/jiv277 (2015).

16 Levine, M., Lu, A., Bennett, D. & Horvath, S. Epigenetic age of the pre-frontal cortex is associated with neuritic plaques, amyloid load, and Alzheimer’s disease related cognitive functioning. Aging (Albany NY) Dec (2015).

17 Levine, M. E. et al. DNA methylation age of blood predicts future onset of lung cancer in the women’s health initiative. Aging (Albany NY) 7, 690–700, doi:10.18632/aging.100809 (2015).

18 Levine, M. E. et al. Menopause accelerates biological aging. Proc Natl Acad Sci U S A 113, 9327–9332, doi:10.1073/pnas.1604558113 (2016).

19 Lu, A. T.et al. DNA methylation GrimAge strongly predicts lifespan and healthspan. Aging (Albany NY) 11, 303–327, doi:10.18632/aging.101684 (2019).

20 Marioni, R. et al. DNA methylation age of blood predicts all-cause mortality in later life. Genome Biol. 16, 25 (2015).

21 Belsky, D. W. et al. Eleven Telomere, Epigenetic Clock, and Biomarker-Composite Quantifications of Biological Aging: Do They Measure the Same Thing? American journal of epidemiology, doi:10.1093/aje/kwx346 (2017).

22 Belsky, D. W. et al. Quantification of the pace of biological aging in humans through a blood test, the DunedinPoAm DNA methylation algorithm. Elife 9, doi:10.7554/eLife.54870 (2020).

23 Olova, N., Simpson, D. J., Marioni, R. E. & Chandra, T. Partial reprogramming induces a steady decline in epigenetic age before loss of somatic identity. Aging Cell 18, e12877, doi:10.1111/acel.12877 (2019).

24 Levine, M. et al. A rat epigenetic clock recapitulates phenotypic aging and co-localizes with heterochromatin. Elife 9, doi:10.7554/eLife.59201 (2020).

25 Petkovich, D. A. et al. Using DNA Methylation Profiling to Evaluate Biological Age and Longevity Interventions. Cell Metab 25, 954–960 e956, doi:10.1016/j.cmet.2017.03.016 (2017).

26 Meer, M. V., Podolskiy, D. I., Tyshkovskiy, A. & Gladyshev, V. N. A whole lifespan mouse multi-tissue DNA methylation clock. Elife 7, doi:10.7554/eLife.40675 (2018).

27 Horvath, S. & Raj, K. DNA methylation-based biomarkers and the epigenetic clock theory of ageing. Nature Reviews Genetics 19, 371–384, doi:10.1038/s41576-018-0004-3 (2018).

28 Liu, Z. et al. Underlying features of epigenetic aging clocks in vivo and in vitro. Aging Cell 19, e13229, doi:10.1111/acel.13229 (2020).

29 Horvath, S. et al. Epigenetic clock for skin and blood cells applied to Hutchinson Gilford Progeria Syndrome and ex vivo studies. Aging (Albany NY) 10, 1758–1775, doi:10.18632/aging.101508 (2018).

30 Lu, A. T. et al. DNA methylation-based estimator of telomere length. Aging (Albany NY) 11, 5895–5923, doi:10.18632/aging.102173 (2019).

31 Hannum, G. et al. Genome-wide methylation profiles reveal quantitative views of human aging rates. Mol Cell. 49, doi:10.1016/j.molcel.2012.10.016 (2013).

32 Zhou, W. et al. DNA methylation loss in late-replicating domains is linked to mitotic cell division. Nat Genet 50, 591–602, doi:10.1038/s41588-018-0073-4 (2018).

33 Yang, Z. et al. Correlation of an epigenetic mitotic clock with cancer risk. Genome Biol 17, 205, doi:10.1186/s13059-016-1064-3 (2016).

34 Teschendorff, A. E. A comparison of epigenetic mitotic-like clocks for cancer risk prediction. Genome Med 12, 56, doi:10.1186/s13073-020-00752-3 (2020).

35 Shireby, G. L. et al. Recalibrating the epigenetic clock: implications for assessing biological age in the human cortex. Brain 143, 3763–3775, doi:10.1093/brain/awaa334 (2020).

36 Chen, B. H. et al. DNA methylation-based measures of biological age: meta-analysis predicting time to death. Aging (Albany NY) 8, 1844–1865, doi:10.18632/aging.101020 (2016).

37 Raj, K. & Horvath, S. Current perspectives on the cellular and molecular features of epigenetic ageing. Exp Biol Med (Maywood) 245, 1532–1542, doi:10.1177/1535370220918329 (2020).

38 Field, A. E. et al. DNA Methylation Clocks in Aging: Categories, Causes, and Consequences. Mol Cell 71, 882–895, doi:10.1016/j.molcel.2018.08.008 (2018).

39 Frick, P. L., Paudel, B. B., Tyson, D. R. & Quaranta, V. Quantifying heterogeneity and dynamics of clonal fitness in response to perturbation. J Cell Physiol 230, 1403–1412, doi:10.1002/jcp.24888 (2015).

40 Blagosklonny, M. V. The hyperfunction theory of aging: three common misconceptions. Oncoscience 8, 103–107, doi:10.18632/oncoscience.545 (2021).

41 Costa-Mattioli, M. & Walter, P. The integrated stress response: From mechanism to disease. Science 368, doi:10.1126/science.aat5314 (2020).

42 Zhang, Y. et al. DNA methylation signatures in peripheral blood strongly predict allcause mortality. Nat Commun 8, 14617, doi:10.1038/ncomms14617 (2017).

43 Kresovich, J. K. et al. Methylation-Based Biological Age and Breast Cancer Risk. J Natl Cancer Inst 111, 1051–1058, doi:10.1093/jnci/djz020 (2019).

44 Babagana, M. et al. Hedgehog dysregulation contributes to tissue-specific inflammaging of resident macrophages. Aging (Albany NY) 13, 19207–19229, doi:10.18632/aging.203422 (2021).

45 Wang, Z. et al. Specific metabolic rates of major organs and tissues across adulthood: evaluation by mechanistic model of resting energy expenditure. Am J Clin Nutr 92, 1369–1377, doi:10.3945/ajcn.2010.29885 (2010).

46 Nathan, A. T. & Singer, M. The oxygen trail: tissue oxygenation. Br Med Bull 55, 96–108, doi:10.1258/0007142991902312 (1999).

47 Sender, R. & Milo, R. The distribution of cellular turnover in the human body. Nat Med 27, 45–48, doi:10.1038/s41591-020-01182-9 (2021).

48 Wang, J. & Zhou, W. H. Epigenetic clocks in the pediatric population: when and why they tick? Chin Med J (Engl) 134, 2901–2910, doi:10.1097/CM9.0000000000001723 (2021).

49 Akdemir, E. S., Huang, A. Y. & Deneen, B. Astrocytogenesis: where, when, and how. F1000Res 9, doi:10.12688/f1000research.22405.1 (2020).

50 Rozhok, A. I. & DeGregori, J. The evolution of lifespan and age-dependent cancer risk. Trends Cancer 2, 552–560, doi:10.1016/j.trecan.2016.09.004 (2016).

51 Higgins-Chen, A. T. et al. A computational solution for bolstering reliability of epigenetic clocks: Implications for clinical trials and longitudinal tracking. bioRxiv, 2021.2004.2016.440205, doi:10.1101/2021.04.16.440205 (2021).

52 Crimmins, E. M., Thyagarajan, B., Levine, M. E., Weir, D. R. & Faul, J. Associations of Age, Sex, Race/Ethnicity, and Education With 13 Epigenetic Clocks in a Nationally Representative U.S. Sample: The Health and Retirement Study. The Journals of Gerontology: Series A 76, 1117–1123, doi:10.1093/gerona/glab016 (2021).

53 Arneson, A. et al. A mammalian methylation array for profiling methylation levels at conserved sequences. bioRxiv, 2021.2001.2007.425637, doi:10.1101/2021.01.07.425637 (2021).

54 Langfelder, P. & Horvath, S. WGCNA: an R package for weighted correlation network analysis. BMC Bioinformatics 9, 559, doi:10.1186/1471-2105-9-559 (2008).

