## Supplemental Figures for "Clock Work: Deconstructing the Epigenetic Clock Signals in Aging, Disease, and Reprogramming"

**Supplementary Figures**

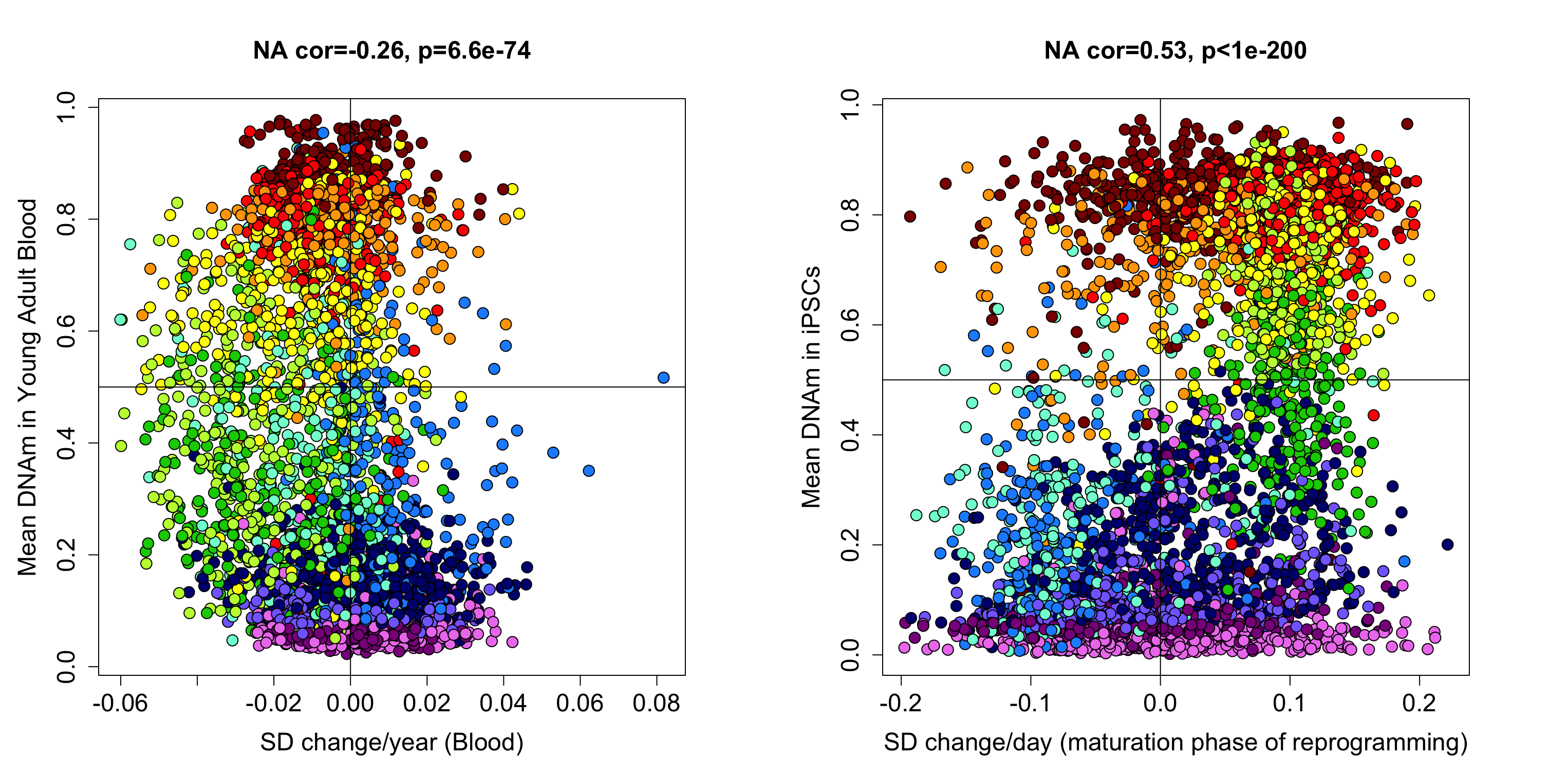

**Figure S1: Standardized changes in DNAm**

For each CpG, we estimated the standardized degree of DNAm change per year in adults by fitting a linear regression model in the Framingham Heart Study (FHS) where standardized betas from CpGs were regressed on age in years. (A) Change per year in standardized DNAm was plotted against the mean DNAm level for each CpG estimated in adults less than 35 years of age from FHS. Point colors denote module assignment, while the vertical line separates CpGs that increase versus decrease with age and the horizontal line distinguishes hypermethylation (DNAm>0.5) from hypomethylation (DNA<0.5). (B) We also plotted standardized changes per day in DNAm during epigenetic reprogramming via Yamanaka expression in dermal fibroblasts against mean DNAm in fully reprogrammed iPSCs. Standardized betas were regressed on days 15-28. Again, point colors denote module assignment, while the vertical line separates CpGs that increase versus decrease with reprogramming and the horizontal line distinguishes hypermethylation (DNAm>0.5) from hypomethylation (DNA<0.5) in iPSCs.

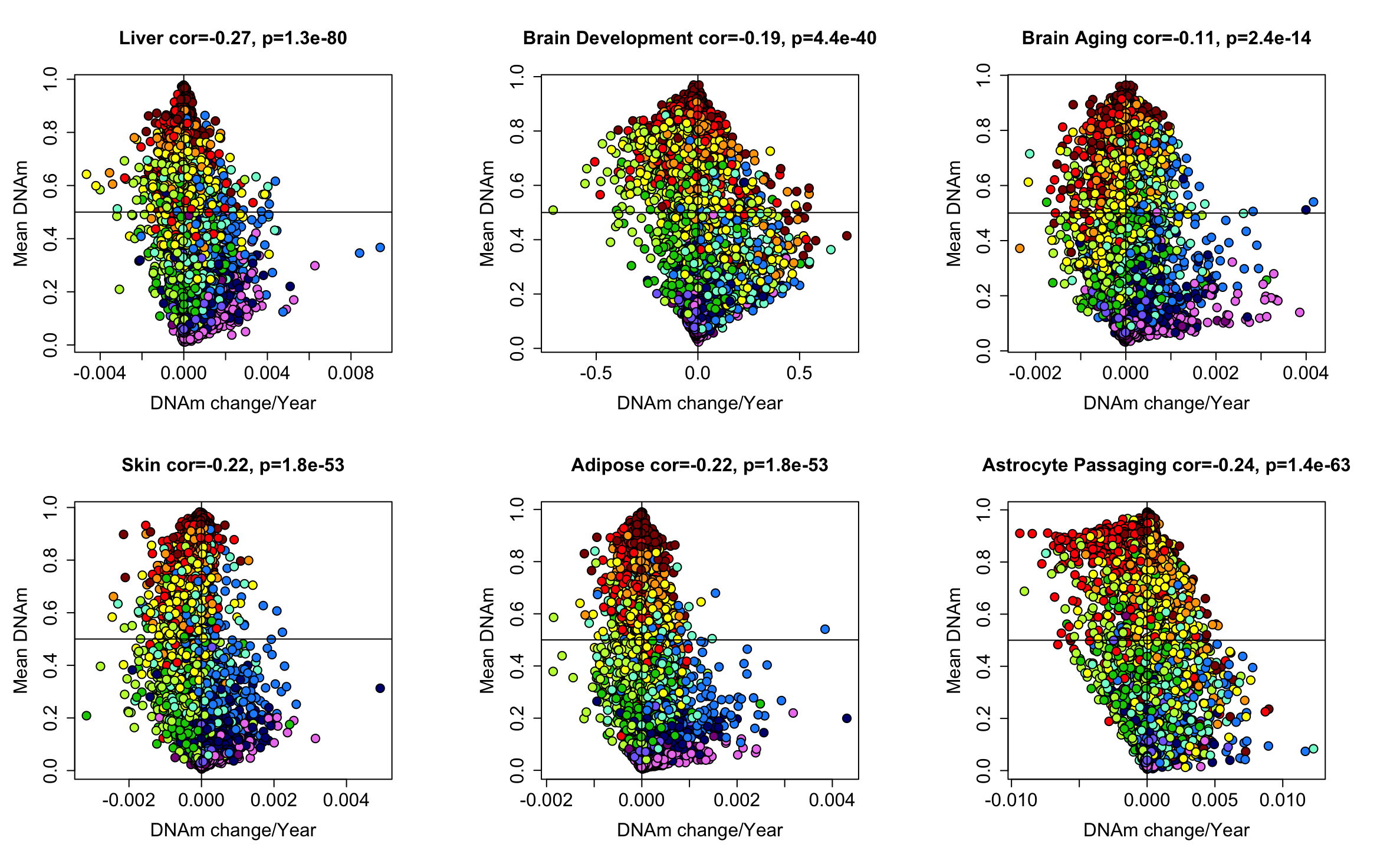

**Figure S2: Module Specific changes in DNAm per year across tissues**

For each CpG, we estimated the sdegree of DNAm change per year in adults by fitting a linear regression model of DNAm betas regressed on age in years for each tissue type—liver, brain (fetal development), brain (adult), skin, adipose. For astrocytes, we estimated change as function of time in culture by regressing DNAm on day. Point colors denote module assignment, while the vertical line separates CpGs that increase versus decrease with age and the horizontal line distinguishes hypermethylation (DNAm>0.5) from hypomethylation (DNA<0.5).

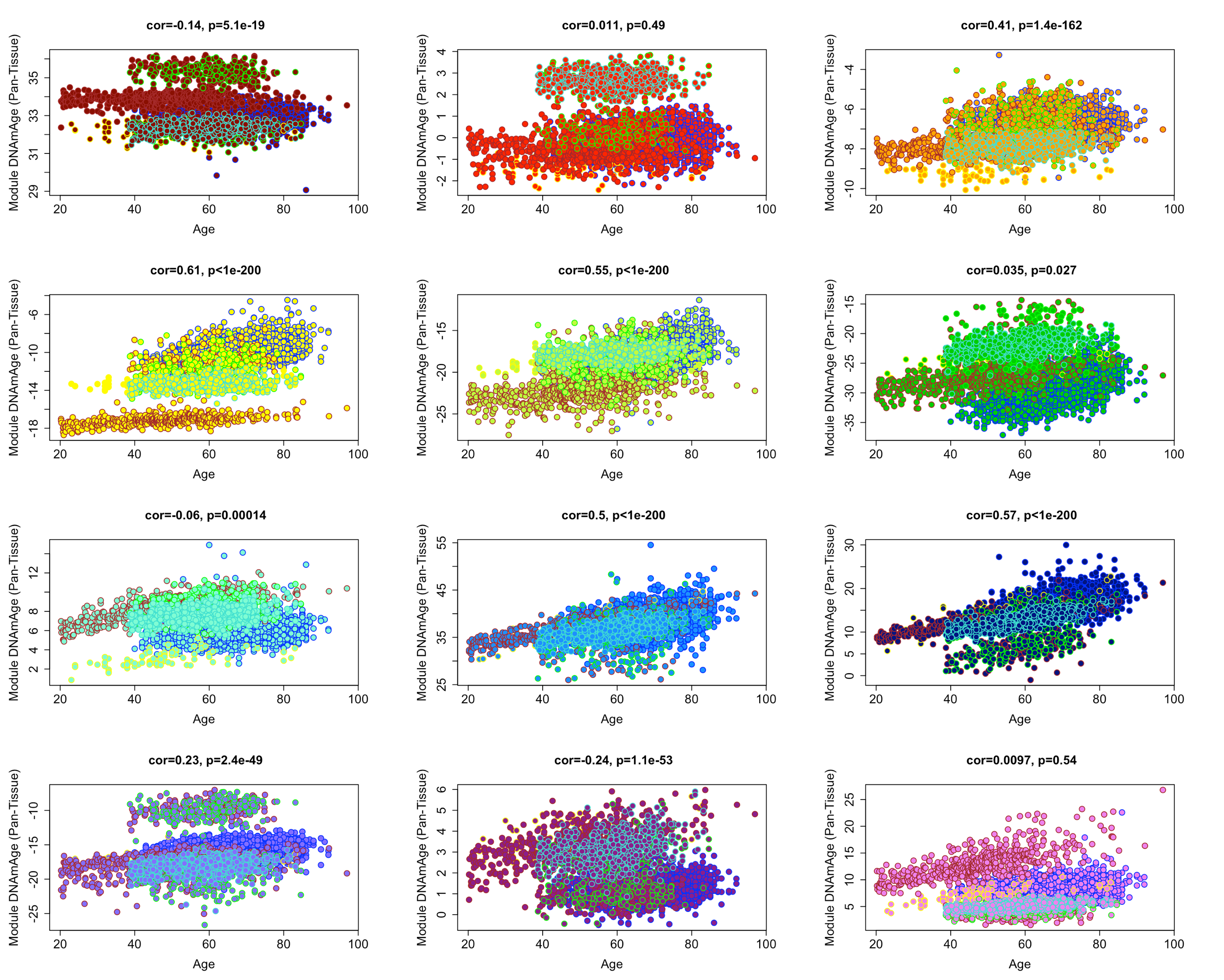

**Figure S3: Multi-tissue age correlations of Pan-Tissue clock modules in adult samples only**

Using the equation for the Horvath Pan-Tissue clock, we estimated module specific DNAmAges and plotted them (y-axis) against chronological age (x-axis) in samples from adult (ages 20+) from blood, liver, skin brain, and adipose. Colors in the upper left and point colors represent the modules. Point outlines represent tissues—brain (brown), blood (blue), skin (green), adipose (turquoise), and liver (yellow).

| EpiTOC2  HypoClock | 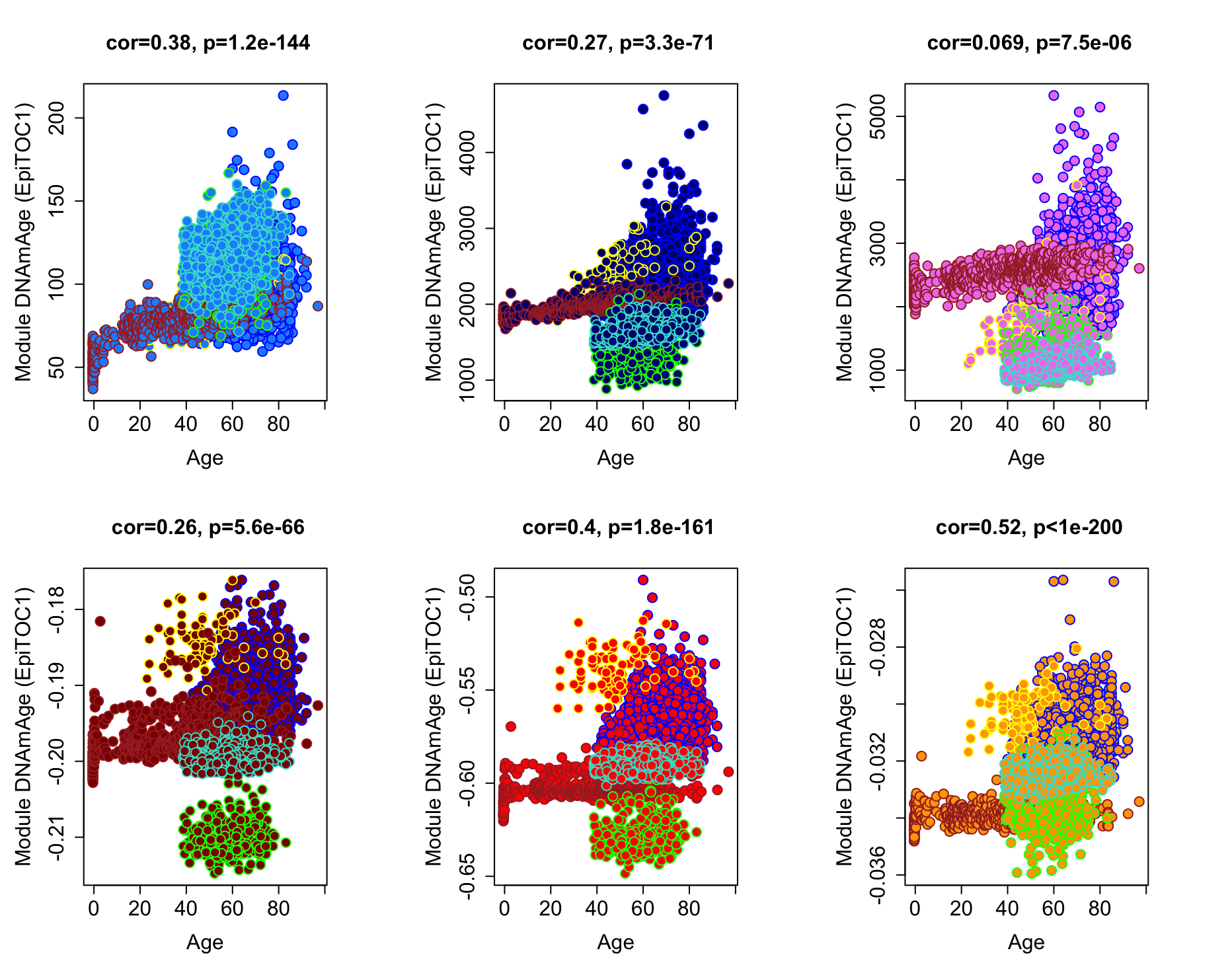 |
| --- | --- |
| **Figure S4: Multi-tissue age correlations of Mitotic Clock modules**  Using the equation for two mitotic clocks—EpiTOC2 and HypoClock—we estimated module specific DNAmAges and plotted them (y-axis) against chronological age (x-axis) in samples from blood, liver, skin brain, and adipose. Colors in the upper left and point colors represent the modules. Point outlines represent tissues—brain (brown), blood (blue), skin (green), adipose (turquoise), and liver (yellow). | |

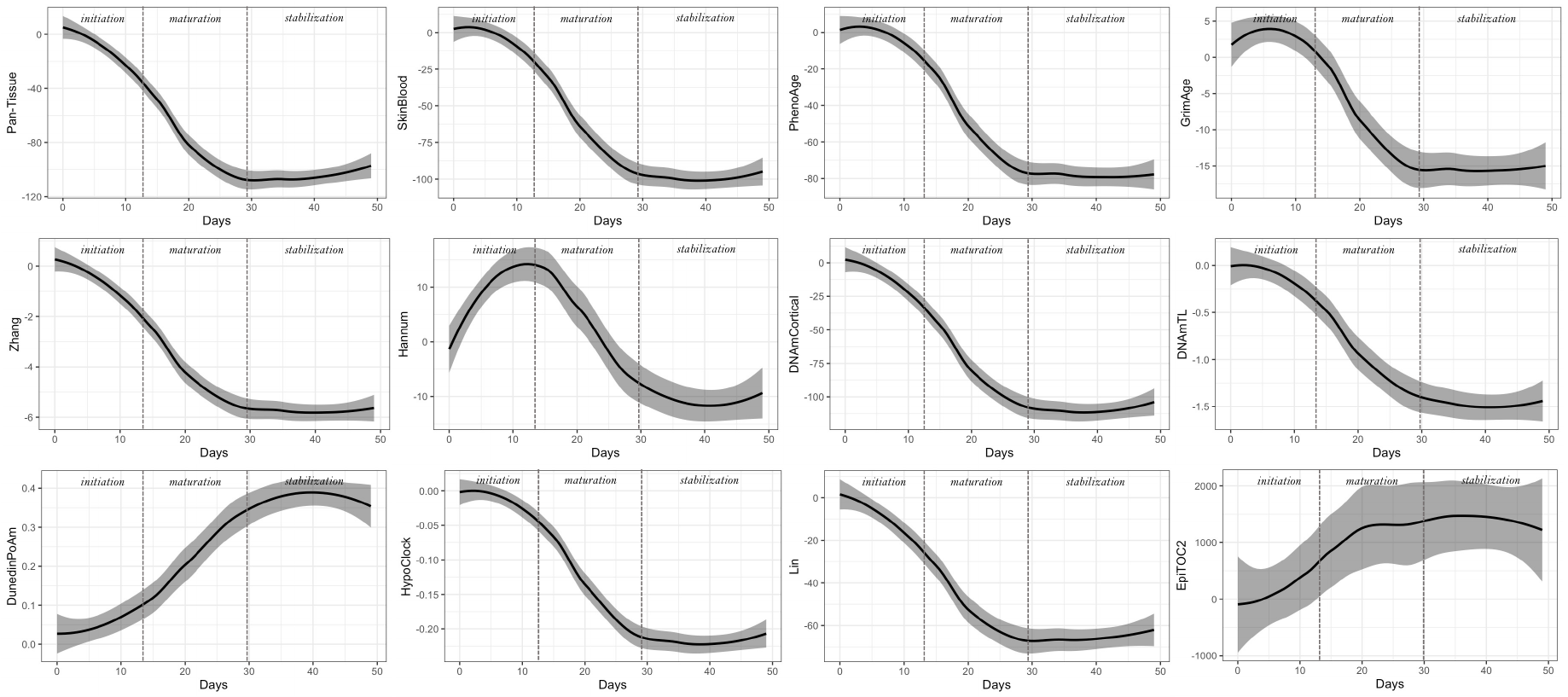

**Figure S5: Time-course effects of fibroblast reprogramming (TRA-1-60 (+) cells) on epigenetic clocks**

DNAmAge were calculated for twelve clocks and plotted as a function of time (days) during reprogramming. We plotted smoothed conditional means (loess) and shaded confidence intervals. Dashed vertical lines depict transitions between initiation, maturation, and stabilization phases. Y-axis depicts the degree of DNAmAge changes since baseline (day 0). We observe strong resetting in most clocks, aside except DunedinPoAm and EpiTOC2, which both showed increases with reprogramming. GrimAge and Hannum did decline in iPSCs, but showed increases at first during the initiation phase, before reverting upon the maturation phase.

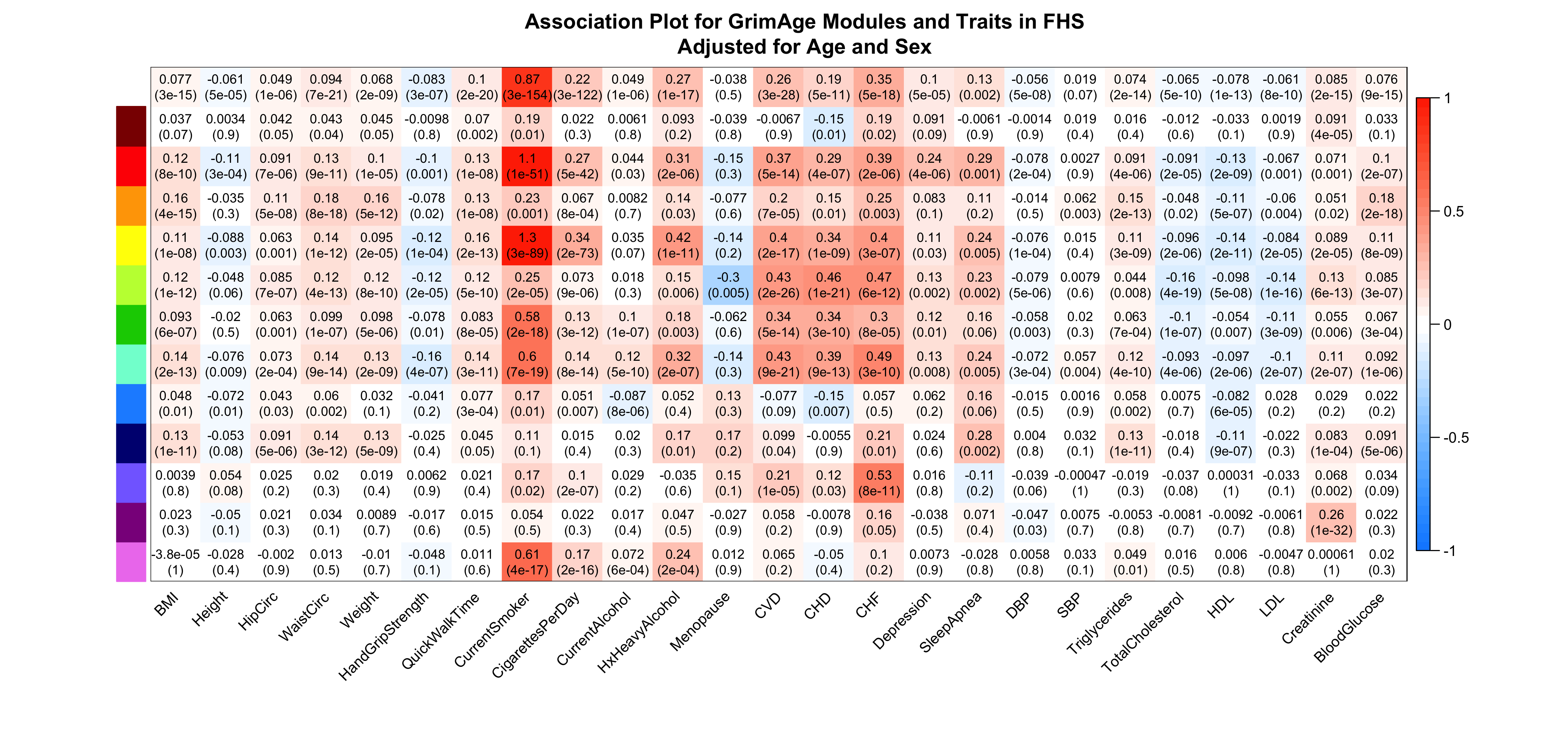

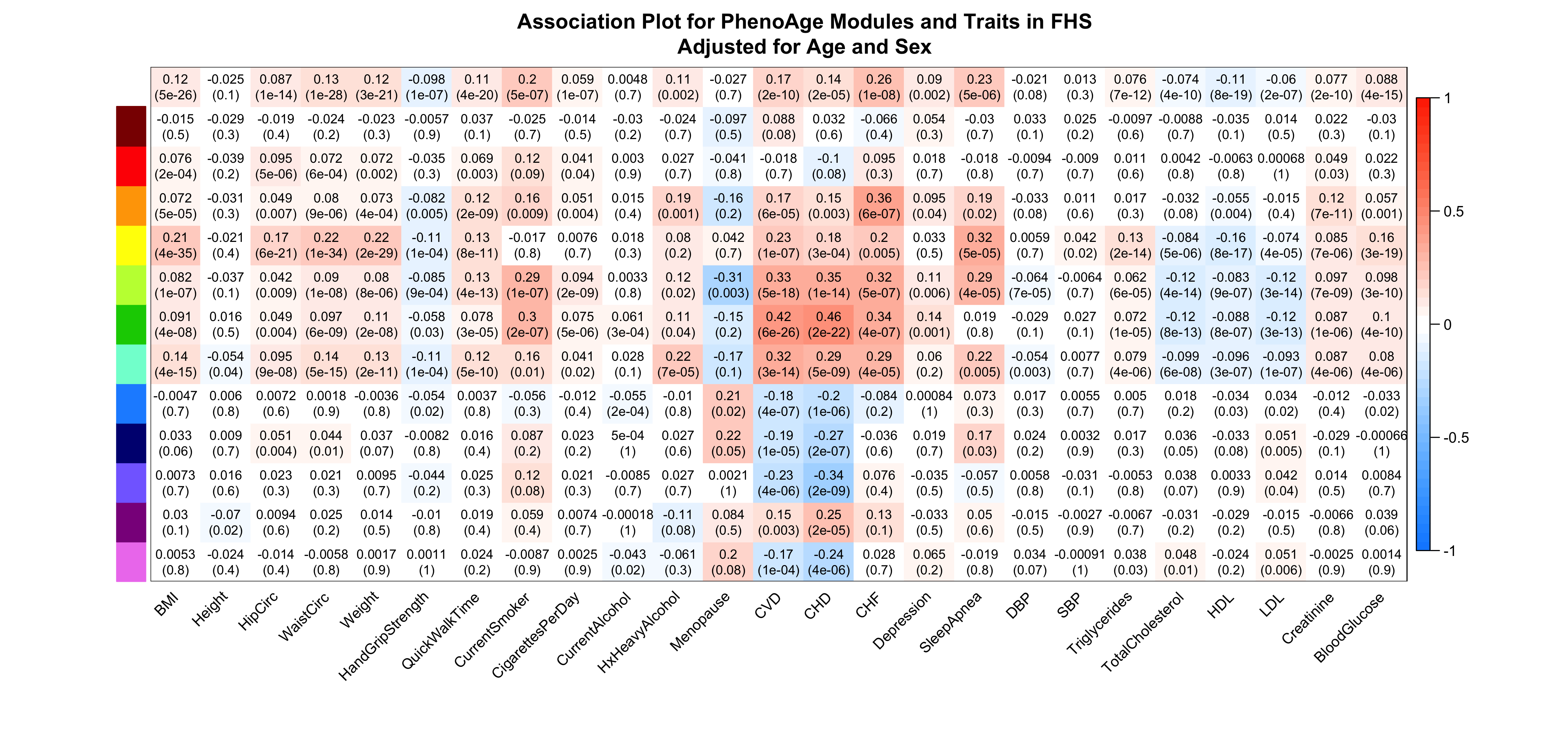

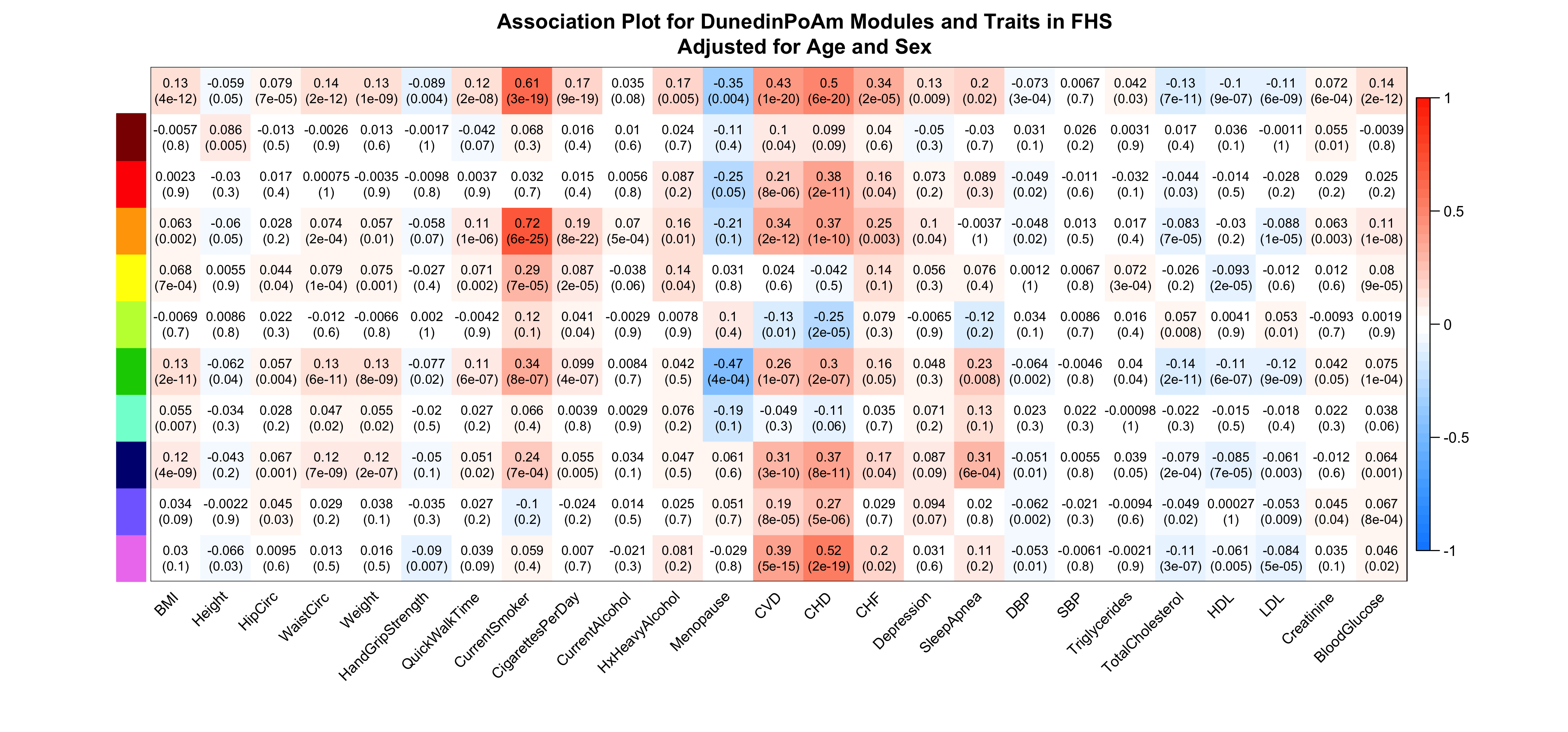

**Figure S6: Associations in blood with modules from three clocks**

We used data from FHS to test for health associations with modules for GrimAge, PhenoAge, and DunedinPoAm. Values depict correlation coefficients (p-values) after residualizing out age and sex. The top row depicts the association for the full clock scores (sum of all modules).

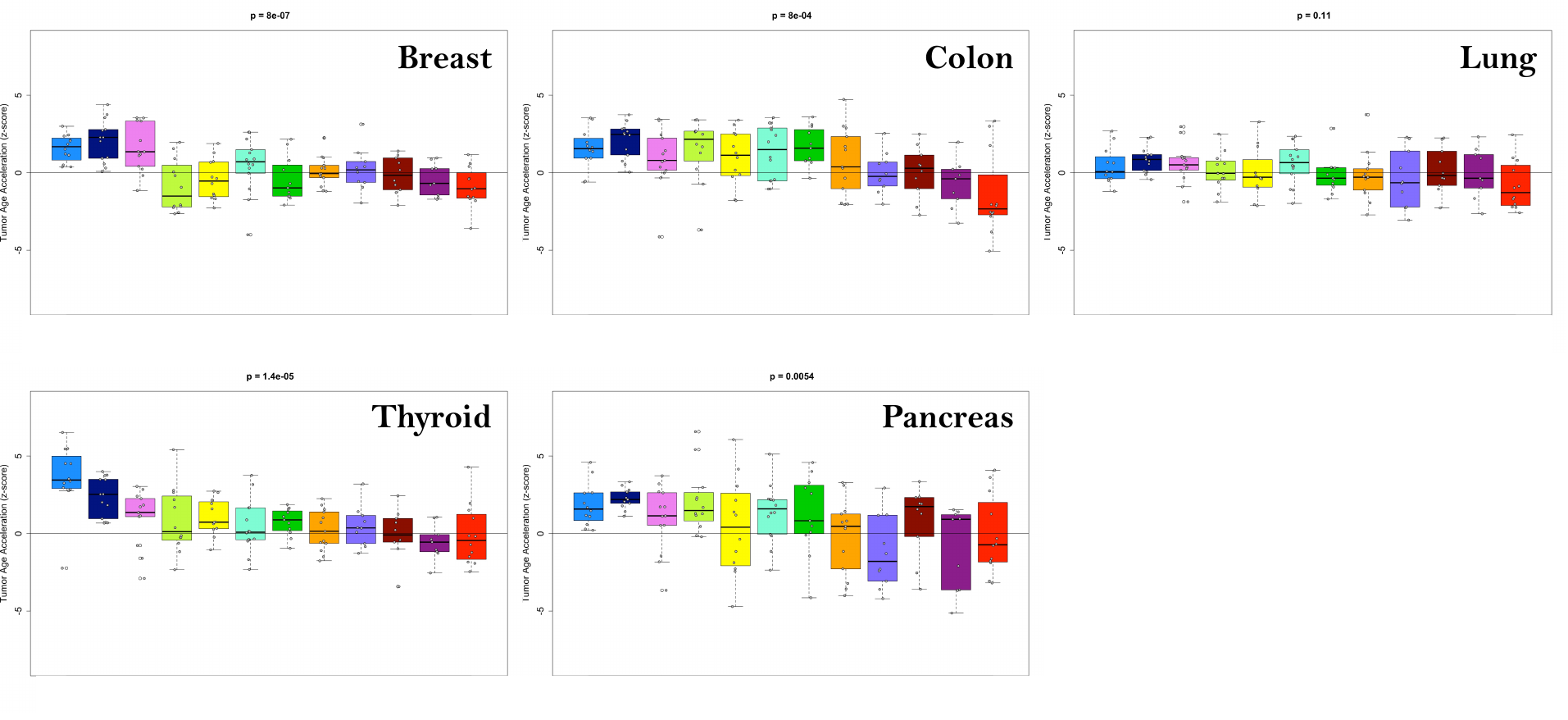

**Figure S6: Tissue specific tumor versus normal tissue differences in clock modules.**

Data from clock modules estimated in tumors and normal tissues, for five tissue types (breast, colon, lung, pancreas, and thyroid) were used to test for accelerated DNAmAge in tumor. The z-scores of effect size (increase in DNAmAge for tumor versus normal tissue) for each clock module pair were extracted from a linear regression model, adjusted for age. Boxplots, with means and standard errors are shown for z-scores pertaining to each module (denoted by color), with positive values suggesting accelerated aging in tumor, and negative values suggesting decelerated aging in tumor. P-value represents significant differences between modules using Kruskal-Wallis test.

\

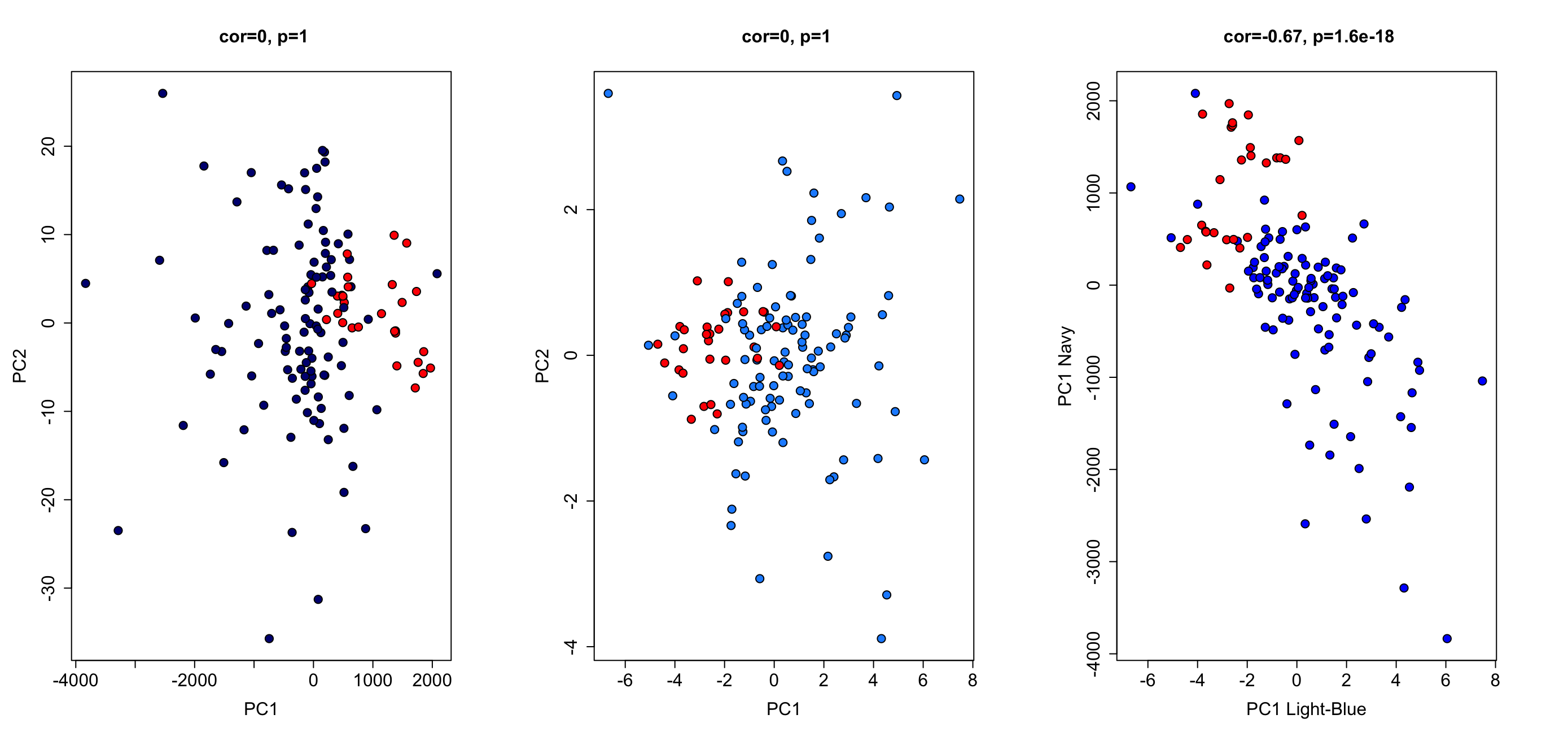

**Figure S7: Navy and light-blue clock modules can mostly differentiate tumor versus normal tissue samples.**

Two PCAs were conducted using navy modules or light-blue modules (adjusted for age and tissue type) from the tumor vs. normal samples as input. We then plotted PC1 versus PC2 for navy where normal samples are shown in red and tumor samples shown in nay. Next we plotted PC1 versus PC2 from light-blue, with normal samples shown in red and tumor samples shown in light-blue. Finally, we plotted PC1 navy versus PC1 light-blue with tumor samples shown in blue and normal shown in red.

**Figure S8: Growth rates and senescence marker accumulation in passaged astrocytes**

**Fetal astrocytes were passaged up to 73 days in culture, with DNAm extracted at each passage. (A)** Cumulative population doubling was calculated using the initial and final cell density, as determined by the countess and plotted against time in culture for 4 biological replicates from two donors. β-gal activity (C12FDG) was measured using flow cytometry or confocal microscopy at each passage and plotted as a function of cPD (B) or passage number (C)
